# TRAF6 controls spinogenesis instructing synapse density and neuronal activity through binding neuroplastin

**DOI:** 10.1101/768341

**Authors:** Sampath Kumar Vemula, Ayse Malci, Lennart Junge, Anne-Christin Lehmann, Ramya Rama, Johannes Hradsky, Ricardo A. Matute, André Weber, Matthias Prigge, Michael Naumann, Michael R. Kreutz, Constanze I. Seidenbecher, Eckart D. Gundelfinger, Rodrigo Herrera-Molina

## Abstract

Synaptogenic mechanisms and their relevance to achieve a correct synapse density and activity in mature neurons are poorly understood. Here, we show that the tumor necrosis factor receptor-associated factor 6 (TRAF6) controls early spinogenesis by binding the cell adhesion molecule neuroplastin which is has been related to synapse formation *in vivo*. TRAF6-neuroplastin co-precipitations from brain samples and co-transfected HEK cells is explained by direct interaction of the proteins based on three-dimensional modelling and biochemical identification of intracellular amino acids of neuroplastin binding the TRAF-C domain of TRAF6 with micromolar affinity. TRAF6 was not only required for normal spinogenesis but also was strictly necessary to restore failed spinogenesis in neuroplastin-deficient neurons. Independently from neuroplastin’s extracellular adhesive properties or interaction with another known partner i.e. the plasma membrane Ca^2+^ ATPases, TRAF6 mediated formation of new postsynapses by neuroplastin overexpression in rat hippocampal neurons. Furthermore, TRAF6-controlled spinogenesis was required for the establishment of a correct synapse density as well as proper synaptic activity and intrinsic neuronal activity as demonstrated with intracellular and extracellular electrophysiological recordings. These findings provide a novel mechanism for early synapse formation that shapes connectivity and functioning of hippocampal neurons.

## Introduction

Synapse formation is a highly coordinated cellular process, which sets up neuronal connectivity in the developing nervous system. Indeed, correct establishment of synapses is fundamental for the information flow in the healthy brain (McAllister 2007; Südhof 2008, 2017) and inaccuracies in synapse formation can underlie altered connectivity in neurological disorders including mental retardation, autism spectrum disorders, and schizophrenia (Südhof 2008, 2017; Zhang et al, 2009; Boda et al, 2010). The appearance of synaptogenic structures in young dendrites, named dendritic protrusions or filopodia, does not seem to be triggered by neuronal activity (Verhage et al., 2000; Sando et al., 2017; Sigler et al. 2017), global intracellular calcium transients (Lohmann et al., 2005; Lohmann and Bonhoeffer 2008) or calcium-dependent signaling (Zhang and Murphy 2004). Rather, it might be controlled by cell-autonomous expression of synaptogenic molecules (Okawa et al., 2014; Jiang et al., 2017a; Südhof 2017). Currently, there is only limited knowledge on how such molecules instruct and organize the formation of synapses. Also, it has not been appreciated how synaptogenic events occurring during the development of neurons contribute to future connectivity or plasticity of the mature brain (Yoshihara et al., 2009; Südhof 2017).

The tumor necrosis factor (TNF) receptor-associated factor 6 TRAF6 is essential for normal brain development. TRAF6 KO embryos display lethal exencephaly and reduced programmed cell death within the developing ventral diencephalon and mesencephalon (Lomaga et al., 2000). Whether TRAF6 plays additional roles in neuronal development for example in synapse formation is unknown. TRAF6 is a largely recognized mediator of inflammatory cell processes, differentiation, activation and tolerance of immune cells as well as morphology and migration of cancer cells (Lomaga et al., 1999; Kobayashi et al., 2001; Xie 2013). TRAF6 is a cytoplasmic E3 ligase and an adaptor protein with an N-terminal region formed by a RING domain and four zinc fingers and a C-terminal region that comprises a coiled coil domain and a TRAF-C domain. The TRAF-C domain is responsible for the binding of TRAF6 to a specific motif in cytoplasmic domains of transmembrane proteins (Chung et al., 2002; Yin et al., 2009). Upon activation, TRAF6 dimers undergo *homo*-oligomerization by lateral engagement of neighboring RING domains leading to a high-order assembly of a three-dimensional lattice-like structure (Yin et al., 2009; Ferrao et al., 2012; Wu 2013). This high-order TRAF6 structures are reported as plasma membrane-associated “fluorescent spots” on the micrometer scale where hundreds of cell signaling intermediaries would nest (Ferrao et al., 2012; Wu 2013).

Neuroplastin is a type-1 transmembrane glycoprotein with a short intracellular tail (Langnaese et al., 1997; Beesley et al., 2014). The isoform neuroplastin 55 (Np55) displays two extracellular immunoglobulin-like (Ig-like) domains whereas neuroplastin 65 (Np65) has an additional N-terminal Ig-like domain with *trans*-homophilic adhesive capacity (Smalla et al., 2000; Owczarek et al., 2011; Herrera-Molina et al., 2014). In humans, neuroplastin has been associated with cortical thickness and cognitive capabilities in adolescents (Desrivieres et al., 2015), schizophrenia (Saito et al., 2007), and Alzheimer’s disease (Ilic et al., 2018). In adult mice, neuroplastin deficiency goes along with retrograde amnesia and impaired associative learning, cortical activity, synaptic plasticity and reduced number of excitatory synapses (Bhattacharya et al., 2017; Herrera-Molina et al. 2014; 2017). Neuroplastin-deficient neurons display lower number of excitatory synapses in the hippocampus (Herrera-Molina et al., 2014; Amuti et al., 2016) and in the inner hear (Carrott et al., 2016; Zeng et al., 2016). Also in hippocampal neurons, inactivation of the *Nptn* gene leads to unbalanced synaptic transmission (Bhattacharya et al., 2017; Herrera-Molina et al., 2014). However, it remains unknown whether neuroplastin participates directly in synapse formation and if this is true, what would it be their specific spinogenic mechanism necessary for the proper establishment of synapse density and whether this would impact synaptic transmission and neuronal activity.

Here, we uncovered a hitherto unanticipated function for TRAF6 in synapse formation. We identified and characterized the TRAF6 spinogenic function as depending on the intracellular binding to neuroplastin, but not in neuroplastin’s extracellular adhesive domain. Also, we demonstrate that TRAF6 binding confers to neuroplastin the cell-autonomous capacity to promote early spinogenesis required directly for the correct establishment of synapse density. Furthermore, we showed that the neuroplastin-TRAF6-controlled spinogenesis is critically relevant for the establishment of a correct synapse density and for the normal synapse activity and adequate functioning of mature neurons.

## Results

### Neuroplastins promote early spinogenesis in young hippocampal neurons

To assess the role of neuroplastin in spinogenesis, we studied the formation of filopodia-like protrusions from dendrites, which can act as precursors of dendritic spines, i.e. postsynaptic compartments of glutamatergic synapses (Ziv and Smith, 1996; McClelland et al., 2010). By confocal microscopy we quantified the number of protrusions per 10 μm length expanding from MAP2-stained dendrites of GFP-filled pyramidal neurons (Figure 1A,B). This indicated that ablation of neuroplastin gene expression results in reduced density of dendritic protrusions in *Nptn^-/-^* compared to *Nptn^+/+^* hippocampal neurons at 9 DIV. The phenotype was rescued by transfection of mutant neurons at 6 or 7 DIV with recombinant neuroplastin isoforms Np55-GFP or Np65-GFP (Figure 1C). In parallel experiments with rat primary hippocampal neurons, we observed that the over-expression of either neuroplastin isoforms promotes dendritic protrusion density (Figure 1D,E). Rat neurons transfected with either Np55-GFP or Np65-GFP at 7 DIV displayed higher density of dendritic protrusion than control GPF-transfected neurons when evaluated at 8 DIV (Figure 1D,E) or 9 DIV (cf. Figure 1F). Surprisingly, later transfections of Np65- or Np55-GFP performed at 9 DIV were ineffective or to raise the protrusion density in rat neurons analyzed at 10 or 11 DIV (Table 1). Therefore, expression of either Np65 or Np55 is mostly effective to raise the density of dendritic protrusions during the early period of spinogenesis, i.e. at 6 to 9 DIV.

**Figure 1.**
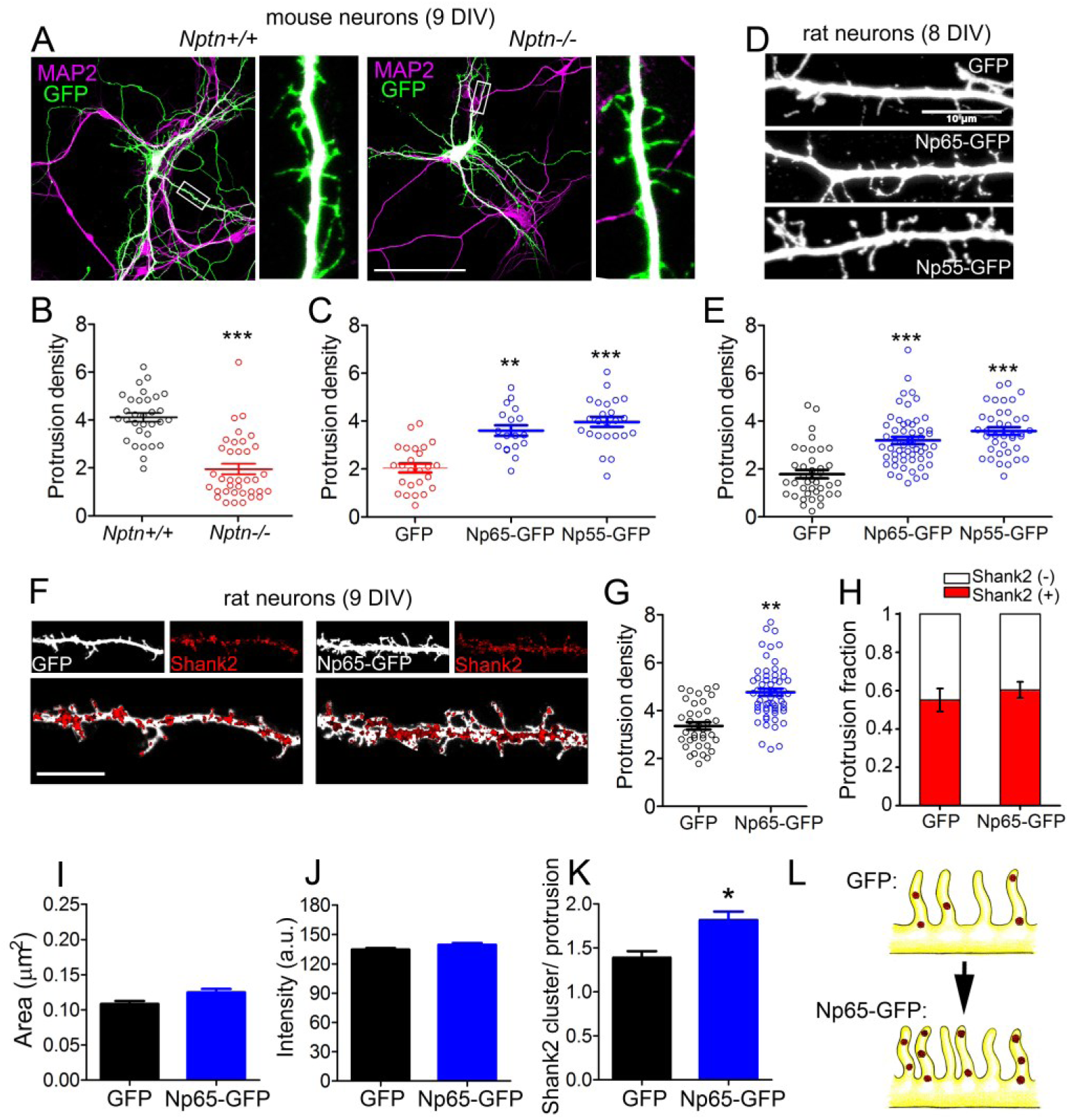
Neuroplastin regulates early spinogenesis. **A**-**C**. Reduced number of dendritic protrusions in *Nptn^-/-^* compared to *Nptn^+/+^* mouse primary hippocampal neurons at 9 DIV. (**A**) *Nptn^-/-^* and *Nptn^+/+^* neurons transfected with GFP-encoding plasmids at 6-7 DIV using Lipofectamine. At 9 DIV, neurons were fixed and stained with anti-GFP antibodies followed by an Alexa 488-conjugated antibody to enhance their intrinsic fluorescence (green) and with anti-MAP2 antibodies followed by a proper secondary antibody to detect dendrites (magenta). Images were obtained using a confocal microscope. Scale bar=100 µm. (**B**) Protrusion density (number of dendritic protrusions per 10μm) of GFP-filled *Nptn^-/-^* and *Nptn^+/+^* neurons (circles) is expressed as mean ± SEM from three independent cultures. ***p<0.001 between genotypes using Student‘s t-test (*Nptn^+/+^* GFP=4.12 ± 0.18 N=33; *Nptn^-/-^* GFP=1.72 ± 0.19 N=36). (**C**) Protrusion density of GFP-, Np65-GFP- or Np55-GFP-expressing *Nptn^-/-^* neurons from two independent cultures. ***p<0.001 or **p<0.01 vs. *Nptn^-/-^* GFP using Student‘s t-test (*Nptn^-/-^* GFP=1.92 ± 0.22 N=26; *Nptn^-/-^* Np65-GFP=3.67 ± 0.18 N=20; *Nptn^-/-^* Np55-GFP=3.77 ± 0.19 N=26). **D**, **E**. Both neuroplastin isoforms increase dendritic protrusion density in rat neurons at 8 DIV. (**D**) Confocal images show rat neurons transfected with plasmids encoding GFP, Np65-GFP or Np55-GFP at 7 DIV. At 8 DIV, neurons were fixed and stained with anti-GFP antibodies followed by an Alexa 488-conjugated antibody (white). Scale bar=10 µm (**E**) Protrusion densities of 40-50 neurons per group (circles) from three-four independent cultures. ***p<0.001 or **p<0.01 vs. GFP transfected cells using Student‘s t-test (GFP=1.95 ± 0.19 N=39; Np65-GFP=3.23 ± 0.14 N=56; Np55-GFP=3.58 ± 0.16 N=38). **F**-**H**. Overexpression of Np65-GFP increases the number of newly formed Shank2-containing spines. (**F**) Confocal images of dendritic segments of rat neurons transfected with GFP or Np65-GFP at 7 DIV. At 9 DIV, neurons were fixed and stained with primary antibodies against GFP (white) and Shank2 (red). Scale bar=10 µm (**G**) Protrusion density (GFP=3.151 ± 0.182 N=48; Np65-GFP=4.642 ± 0.145 N=54) and (**H**) Distribution of Shank2-positve and Shank2-negative protrusions were calculated as a fraction from 40-50 neurons per group from three independent experiments. Plots display mean ± SEM as indicated. **p<0.01 for Np65-GFP vs GFP using Student‘s t-test (Shank2(+): GFP=0.54 ± 0.07; Np65-GFP=0.60 ± 0.06). **I**-**K**. (**I**) Size of puncta (area; GFP=0.10 ± 0.01 N=747; Np65-GFP=0.11 ± 0.01 N=738), (**J**) Fluorescence intensity (GFP=127.6 ± 2.1; Np65-GFP=131.5 ± 1.9) and (**K**) Number of Shank2 clusters/protrusion in neurons (GFP=1.46 ± 0.17 N=43; Np65-GFP=1.91 ± 0.18 N=49) of the experiments displayed in Figure 1F. *p<0.05 between Np65-GFP-expressing and GFP-expressing neurons using Student‘s t-test. **L**. The upper sketch on the left illustrates dendritic protrusions enriched on Shank2 in control GFP-filled hippocampal neurons at 9 DIV. Np65-GFP-expressing neurons (lower sketch) display more spinogenic protrusions with Shank2 clusters.

**Table 1.**
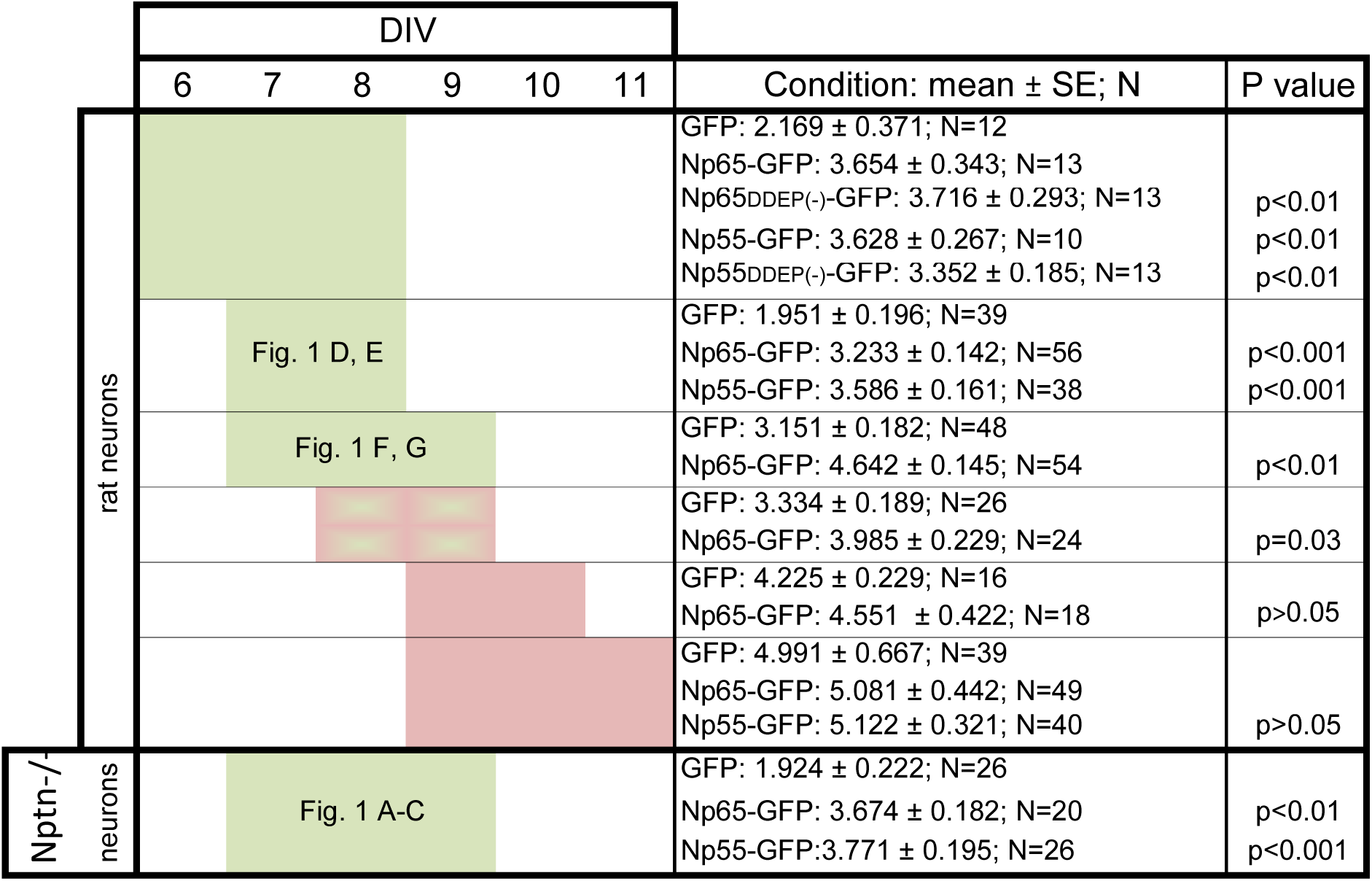
Time course of Np-controlled spinogenesis. The table is explained in the main text and in Figure 1. Briefly, time periods when Np mediates spinogenesis around 6 to 9 DIV are highlighted in green.

To characterize the nature of neuroplastin-promoted dendritic protrusions, we evaluated the appearance of Shank2, a well-established marker (Grabrucker et al., 2011; Sarowar and Grabrucker 2016) and key organizer (Roussignol et al., 2005) of functional postsynapses. While transfection of Np65-GFP at 7DIV increased dendritic protrusion density compared to transfection with control GFP, the relative abundance of Shank2-positive vs. Shank2-negative protrusions (Protrusion fraction) was not different between Np65-GFP-tranfected and GFP-transfected rat neurons at 9 DIV (Figure 1F,H). Although area and intensity of Shank2 clusters were not different between Np65-GFP- and GFP-expressing dendrites at 9 DIV (Figure 1I,J), the number of Shank2 clusters per dendritic protrusion was higher in Np65-GFP-vs. GFP-expressing dendrites at 9 DIV (Figure 1K). The data show that Np65-GFP-overexpressing neurons display an increased number of newly formed postsynapses defined as dendritic protrusions containing a higher number of Shank2 clusters compared to GPF-filled control neurons (Figure 1L). These results are consistent with the idea that both neuroplastin isoforms share the potential to promote the formation of dendritic protrusion during a time period of spinogenesis. Accordingly, we looked for additional *cis*-acting mechanisms related to both neuroplastins to increase the density of spinogenic dendritic protrusions.

### A TRAF6 binding motif is present in neuroplastin, but not in others synaptogenic CAMs

To decipher underlying signaling mechanisms of spinogenesis, we sought for potential intracellular binding partners of neuroplastins employing *in silico* approaches. Using the ELM database (http://elm.eu.org/) we identified a single TRAF6 binding motif in the cytoplasmic tail of all neuroplastins from human, rat, and mouse (Figure 2A) which fully matches the well-characterized TRAF6 binding motif (Ye at al., 2002; Sorrentino et al., 2008; Yin et al., 2009). Very surprisingly, the TRAF6 binding motif was not found amount a number of other known spinogenic type-1 CAMs namely N-Cadherin (Bozdagi et al, 2010), LRRTM (Linhoff et al, 2009), neuroligins (Varoqueaux et al, 2006), neurexins (Missler et al; 2003), SynCAM1 (Robbins et al, 2010), EphB2 (Henderson et al, 2001), PTPR0 (Jiang et al., 2017), and others (Figure S1). These results highlight both the specificity and novelty of the association of TRAF6 to neuroplastin to mediate spinogenesis. After further analysis, we noticed that the alternative mini-exon-encoded Asp-Asp-Glu-Pro (DDEP) of neuroplastin is close to its TRAF6 binding motif (Figure 2A). Based on crystallographic studies on the interaction of the TRAF6 TRAF-C domain with the TRANCE receptor (Ye et al., 2002), *in silico* modelling was applied to TRAF6 TRAF-C domain-neuroplastin interaction (Figure 2B; Figure S2A). A strikingly similar three-dimensional structure was predicted for the TRAF6 binding site of neuroplastin when compared to the TRANCE receptor TRAF6 binding motif (Figure 2B). In particular, the coordinates and stereo specificity of key amino acids (Figure 2B: P_-2_= Pro, P_0_= Glu, and P_3_= Aromatic/Acidic) involved in docking of the TRANCE receptor to TRAF6 TRAF-C domain (TRAF-C) were conserved in the TRAF6 binding site of neuroplastin (Figure 2C). Thus, we conclude that the cytoplasmic tail of neuroplastins displays a proper TRAF6 binding site.

**Figure 2.**
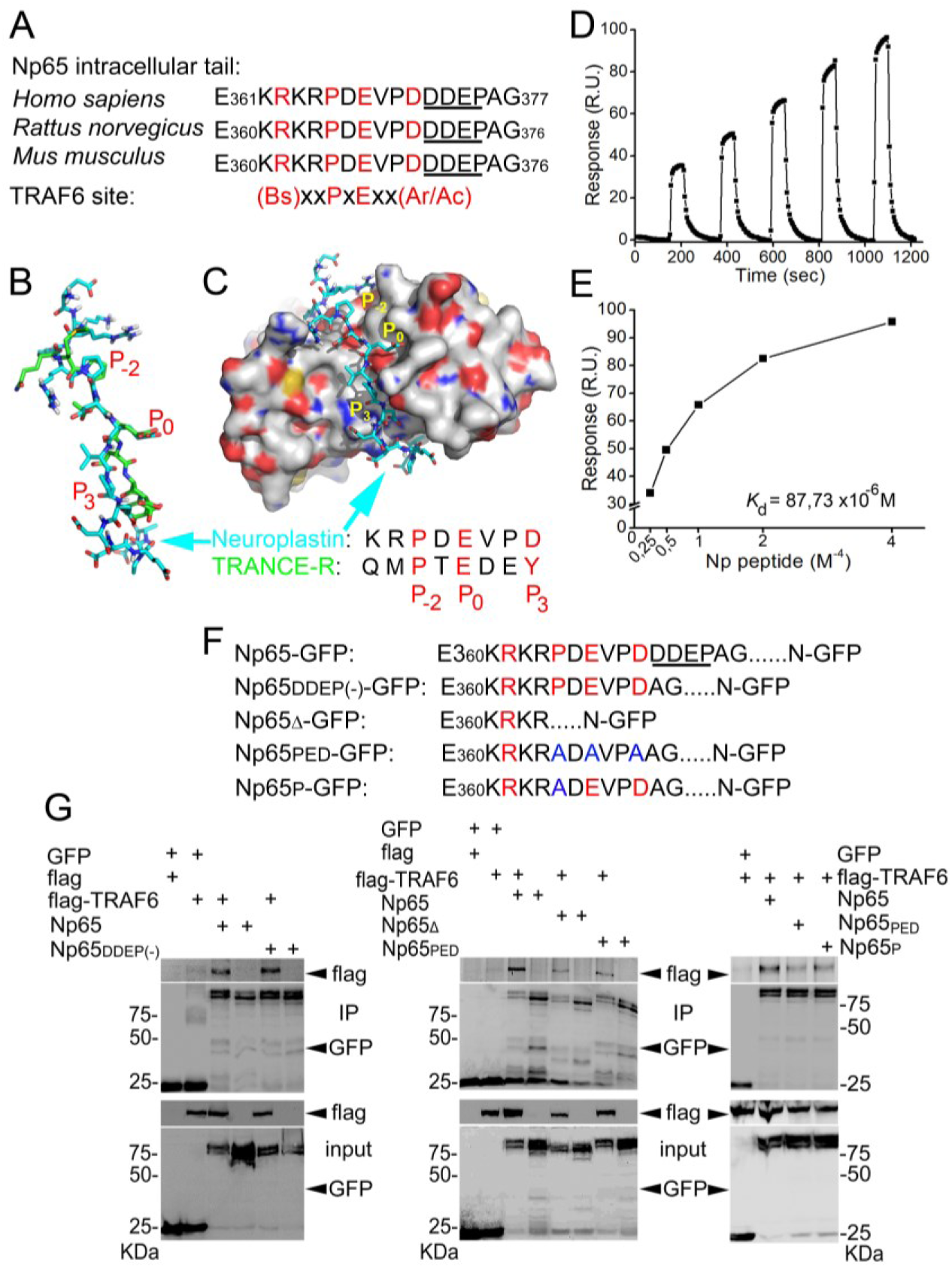
Characterization of the binding of TRAF6 to neuroplastin. **A**. Potential TRAF6 binding site in the intracellular tail of neuroplastin 65 (and identical in neuroplastin 55) fits the canonical and specific motif recognized by TRAF6. The alternatively spliced DDEP sequence is underlined. Bs, Ac, and Ar stand for basic, acidic, and aromatic amino acids, respectively. **B**, **C**. Neuroplastin-TRAF6 binding *in silico*. (**B**) Three-dimensional model of the TRAF6 binding site in the intracellular tail of neuroplastin (cyan) and key amino acids responsible for the binding to TRAF6 fit to the well-known TRAF6 binding motif present in the TRANCE receptor (green). (**C**) Docking of the TRAF6 C-domain with the TRAF6 binding site of neuroplastin. Similar to the binding of TRANCE receptor to TRAF6 documented by a crystallographic study (Yin et al., 2009), interaction of neuroplastin with TRAF6 would be mediated by the Proline (P) in the position P_-2_ Glutamic acid (E) in P0, and Aspartic acid (D) in P_3_. **D**, **E**. Direct binding of the neuroplastin-derived intracellular peptide comprising the TRAF6 binding site to purified recombinant TRAF6. The binding curve (**D**) and the binding curve (**E**) for the neuroplastin-TRAF6 binding where obtained using surface plasmon resonance. **F**, **G**. Neuroplastin-TRAF6 co-precipitation is drastically decreased by deletion or mutation of key amino acids in the TRAF6 binding site of neuroplastin. (**F**) Neuroplastin constructs included into the experiments. (**G**) HEK cells were co-transfected with constructs encoding either GFP, Np65-GFP or Np65_DDEP(-)_-GFP and with TRAF6-flag or flag alone for 24 hours. Alternatively, cells were co-transfected with GFP, Np65-GFP, Np65Δ-GFP (TRAF6 binding site deficient construct), Np65_PED_-GFP (containing a TRAF6 binding site with triple substitution to alanine) or Np65_P_-GFP (with single substitution to alanine) and with TRAF6-flag or flag constructs for 24 hours. After homogenization, anti-GFP antibody-coupled beads were used to precipitate GFP-tagged complexes. We used anti-Flag or anti-GFP antibodies to detect the proteins as indicated.

### TRAF6 binds neuroplastins

We confirmed that TRAF6 co-precipitates with neuroplastins from brain homogenates (Figure S2E). Based on this result, we decided to test whether there is a direct interaction between TRAF6 and neuroplastins. Thus, we characterized the binding of purified neuroplastin intracellular peptide containing the TRAF6 binding motif to immobilized recombinant TRAF6 by surface plasmon resonance (Figure 2D,E; Figure S2B,C). Binding of neuroplastin peptide to TRAF6 was dependent on peptide concentration, saturable, and displayed a 1:1 stoichiometry. We calculated a K_d_ value of 88 μM for the neuroplastin-TRAF6 interaction (Figure 2D,E), which is very similar to the K_d_ of 84 μM for the TRANCE receptor-TRAF6 interaction (Yin et al., 2009). To establish whether the TRAF6 motif in neuroplastin binds TRAF6 in living cells, we performed co-immunoprecipitation assays from HEK cells transfected with different GFP-tagged constructs of neuroplastins and flag-tagged TRAF6. Due to alternative splicing of the primary transcript both major neuroplastin isoforms can contain an alternative DDEP insert close to their TRAF6 binding site. To consider potential differences in binding, splicing variants with and without DDEP were tested. As shown in Figure 2F,G both splice variants of Np65-GFP co-precipitated flag-TRAF6 suggesting that the mini exon-encoded insertion is not critical for the interaction. Similarly, Np55 with and without DDEP insertion co-precipitated with TRAF6 (Figure S2D). In contrast, co-precipitation was largely abolished when GFP-tagged versions of Np65 either with deleted TRAF6 binding motif (Np65Δ-GFP) or with triple (Np65_PED_-GFP) or single (Np65_P_-GFP) amino acid substitutions in the binding site (Figure 2F-G) were used. Additionally, pull-down assays demonstrate that Np65-GFP isolated from HEK cell homogenates binds equally well to purified recombinant GST-TRAF6 or to GST-TRAF6 C-domain (coiled coil and TRAF-C domains GST-TRAF6_cc-c_) (Figure EV1B,C). Based on these analyses, we conclude that the TRAF6 binding site in the cytoplasmic tail of neuroplastins is fully capable of binding the TRAF-C domain of TRAF6.

### TRAF6 mediates the formation of filopodial structures by neuroplastin

We have reported disorganization of polymerized actin in dendrites of *Nptn^-/-^* primary hippocampal neurons (Herrera-Molina et al., 2014). Coincidently, TRAF6 has been associated to regulation of actin polymerization (Armstrong et al., 2002; Wang et al., 2006; Yamashita et al., 2008). As a first test to understand the significance of the neuroplastin-TRAF6 interaction, we therefore performed experiments to explore if and how neuroplastins and TRAF6 interact to regulate actin-based filopodia formation in HEK cells. Over-expression of Np65 or Np55 in HEK cells was sufficient to induce a massive increase of filopodia number and length as compared to control cells transfected with either soluble or membrane-attached GFP (Figure 3A-C). Variants of Np55 or Np65 lacking the DDEP insert were as effective as the ones that carry the insert to promote filopodial structures (Figure S3A-D). The capacity of neuroplastin to promote filopodia was eliminated by mutants abolishing TRAF6 binding (i.e. Np65Δ-GFP, Np65_PED_-GFP, Np65_P_-GFP) (Figures 3A-C). Furthermore, after decreasing protein levels of endogenous TRAF6 by ∼80% using a specific siRNA (Figure S3E,F), neither expression of Np65-GFP nor of Np55-GFP did increase the number or length of filopodia in HEK cells (Figure 3A-C). Thus, Np55 and Np65 (±DDEP) are equally effective to promote the formation of filopodial structures and they seem to require endogenous TRAF6 and binding to their TRAF6 motifs to do so.

**Figure 3.**
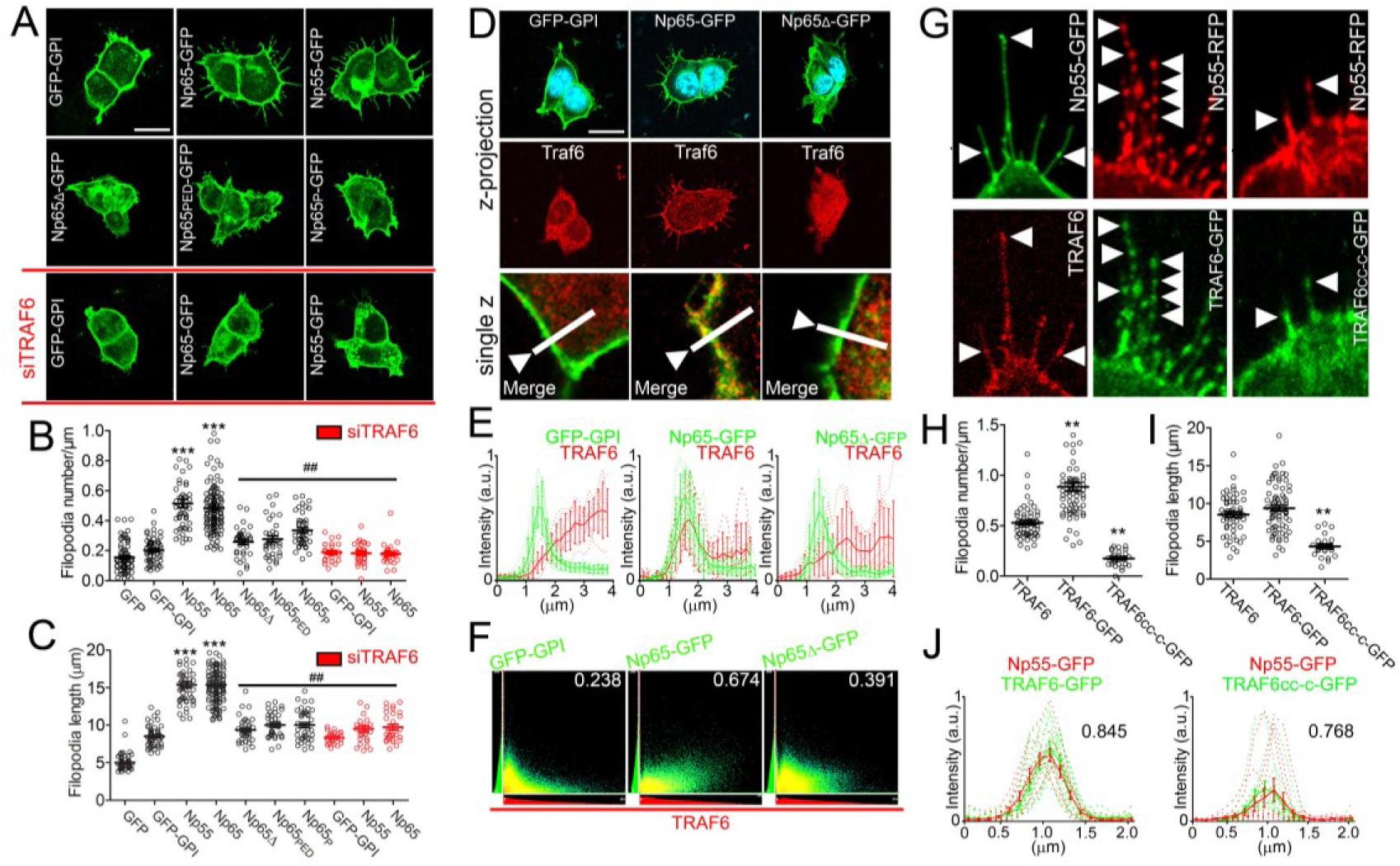
Neuroplastin requires its TRAF6 binding site and TRAF6 to promote filopodia formation. For these experiments, HEK cells were transfected with plasmids coding for either soluble GFP, membrane-attached GFP-GPI, DDEP insert-containing isoforms Np65-GFP or Np55-GFP, TRAF6 binding site-deficient Np65Δ-GFP, Np65_PED_-GFP (containing triple substitution to alanine in the TRAF6 binding site) or Np65_P_-GFP (with single substitution to alanine), full-length TRAF6-GFP or coiled coil-TRAF-C-GFP (TRAF6_cc-c_-GFP). To knockdown endogenous TRAF6, cells were transfected with siRNA against TRAF6 or scrambled siRNA for 24 hours before the transfection with plasmids. After 24 hours, cells were fixed with methanol and immunostained with anti-GFP rabbit antibody overnight and with an Alexa-488 secondary antibody. Alternatively, cells were additionally stained with an anti-TRAF6 rabbit antibody followed by a Cy3-conjugated secondary antibody and DAPI. **A**-**C**. Deletion or mutations of the TRAF6 binding site of neuroplastin or knockdown of endogenous TRAF6 decrease neuroplastin capacity to promote filopodia. (**A**) Scale bar=10 µm. (**B**) Filopodia number per micron of plasma membrane (GFP=0.15 ± 0.01 N=70; GFP-GPI=0.20 ± 0.01 N=59; Np55-GFP=0.51 ± 0.02 N=51; Np65-GFP=0.48 ± 0.01 N=126; Np65Δ-GFP=0.25 ± 0.01 N=34; Np65_PED_-GFP=0.27 ± 0.02 N=34; Np65_P_-GFP=0.33 ± 0.01 N=40; siTRAF6 GFP-GPI=0.29 ± 0.01 N=30; siTRAF6 Np55-GFP=0.17 ± 0.01 N=28; siTRAF6 Np65-GFP=0.18 ± 0.01 N=27) and (**C**) Filopodia length (GFP=5.07 ± 0.24; GFP GPI=8.41 ± 0.33; Np55-GFP=16.68 ± 0.76; Np65-GFP=16.67 ± 0.48; Np65Δ-GFP=9.48 ± 0.57; Np65_PED_-GFP= 10.79 ± 0.58; Np65_P_-GFP=10.78 ± 0.64; siTRAF6 GFP-GPI= 1.73 ± 0.72; siTRAF6 Np55-GFP=10.18 ± 0.72; siTRAF6 Np65-GFP= 9.77 ± 0.85) were quantified using a semi-automatized Matlab-based algorithm. ***p<0.001 for the indicated condition vs. GFP and ^##^p<0.01 vs. Np65 using Student‘s t-test. **D**-**F**. Full-length neuroplastin recruits cytosolic TRAF6 to the cell membrane. (**D**) Endogenous TRAF6 is recruited by and co-localizes with Np65-GFP, but not with Np65Δ-GFP nor with GFP-GPI at the plasma membrane. Scale bar=10 µm. The lower confocal pictures are single z plains and on them, a line scan served to quantify the fluorescence distribution of the GFP-tagged proteins and TRAF6 as shown in (**E**). (**F**) Co-localization index (Pearson’s coefficient) is displayed for each of the condition as indicated. **G**-**J**. Elimination of RING domain abrogates TRAF6 capacity to mediate neuroplastin-promoted filopodia formation. (**G**) The pictures are single z plains acquired by confocal microscopy. Arrow heads point to fluorescent spots formed by Np55-RFP co-localizing with TRAF6-GFP or with TRAF6_cc-c_-GFP. (**H**) Filopodia number (TRAF6=0.53 ± 0.02 N=54; TRAF6-GFP=0.88 ± 0.04 N=69; TRAF6_cc-c_-GFP=0.17 ± 0.02 N=25) and (**I**) Filopodia length (TRAF6=13.44 ± 0.35; TRAF6GFP=13.89 ± 0.37; TRAF6_cc-c_-GFP= 4.3 ± 0.29) were obtained using a semi-automatized Matlab-based algorithm. **p<0.01 vs. TRAF6 using Student‘s t-test. (**J**) Line scan analysis of TRAF6-GFP- and Np55-RFP-associated fluorescent signals of single spots. Also, the corresponding co-localization index (Pearson’s coefficient) is displayed for each plotting.

TRAF6 translocates from the cytoplasm to the membrane by recruitment to integral membrane proteins with TRAF6 binding domains (Yin et al., 2009; Wu 2013). Therefore, we tested whether neuroplastins *via* their C-terminal TRAF6 binding motif have the capacity to recruit endogenous TRAF6 to the plasma membrane. In HEK cells transfected with GPI-anchored GFP or with Np65Δ-GFP, TRAF6 immunoreactivity was primarily located in the cytoplasm (Figure 3D). In contrast, TRAF6 immunoreactivity was abundantly associated with the plasma membrane in cells expressing recombinant Np65-GFP (Figure 3D) or other variants of neuroplastin (Figure S3A). Analyses of co-distribution (Figure 3E) and co-localization (Figure 3F) confirmed that plasma membrane-associated TRAF6 co-localizes with Np65. Thus, neuroplastin can recruit TRAF6 to the plasma membrane and thereby regulate its subcellular localization. This is independent of the presence or absence of the DDEP insert.

Next, we asked whether the recruitment and binding of TRAF6 by neuroplastin mediate filopodia formation. To test this prediction, we co-expressed GFP-tagged TRAF6 (TRAF6-GFP) with Np55-RFP. Clearly, co-expression of TRAF6-GFP fostered the increase of filopodia number by Np55-GFP (Figure 3G-I). Very interestingly, endogenous TRAF6 and TRAF6-GFP co-localized with Np55-RFP in filopodia-associated microscopic spots (Figure 3G). Indeed, analyses of fluorescent intensity and distribution revealed high co-localization of TRAF6-GFP with Np55-RFP in single spots of filopodia (Figure 3J). The potential involvement of the N-terminal RING domain of TRAF6 was tested using TRAF6_cc-c_-GFP containing the coiled coil and TRAF-C domains and lacking the N-terminal domain (see Figure S2B,C). Despite being recruited to the plasma membrane and co-localized with Np55-RFP (Figure 3G,J), TRAF6_cc-c_-GFP blocked neuroplastin-induced filopodia formation (Figure 3G-I). Accordingly, the recruitment and binding of TRAF6_cc-c_ by neuroplastin is insufficient to promote filopodial structures. Because RING domain is well-known to be responsible for three-dimensional assembly of functional TRAF6 lattice-like structures (Yin et al., 2009; Ferrao et al., 2012; Wu 2013), we conclude that only the recruitment and binding of fully functional TRAF6 can mediate formation of filopodial structures by neuroplastin.

### TRAF6 confers spinogenetic capacity to neuroplastin

We tested the hypothesis that TRAF6 is essential for the spinogenic function of neuroplastin in hippocampal pyramidal neurons. As initial evidence for this, we confirmed the co-localization/co-distribution of TRAF6 and neuroplastin using high-resolution microscopy and image deconvolution procedures in dendritic protrusions of young neurons but not in spines of mature neurons (Figure S4). In addition, we evaluated the density of dendritic protrusions after decreasing levels of TRAF6 by siRNA knockdown (as characterized in Figure S3E,F) in rat neurons transfected with GFP-, Np65-GFP or Np65Δ-GFP at 9 DIV. When TRAF6 levels were knocked down by 60% or more, the dendritic protrusion density was reduced in GFP-, Np65-GFP-, and Np65Δ-GFP-expressing neurons (Figure S3C,D).

Then, we used *Nptn^-/-^* hippocampal neurons with significantly reduced number of spinogenic protrusions (Figure 1). The protrusion density in *Nptn^-/-^* dendrites expressing Np65-GFP was higher than in control *Nptn^-/-^* dendrites expressing GFP at 9 DIV (Figure 4A,B). Clearly, Np65Δ-GFP failed to rescue the dendrite protrusion density in *Nptn^-/-^* neurons (Figure 4A,B). In independent experiments, we found that Np65Δ-GFP-failed to promote the density of dendritic protrusions as Np65Δ-GFP-expressing rat neurons displayed a similar density of dendritic protrusions as GPF-expressing rat neurons at 8 DIV (Figure 4C,D). Additionally, we evaluated whether Np65Δ-GFP affects the number of protrusions and interferes with the normal enrichment of Shank2 in dendritic protrusions in rat neurons at 9 DIV. The density of dendritic protrusions and the distribution of Shank2-positive vs. Shank2-negative protrusions were similar between GFP- and Np65Δ-GFP-expressing rat neurons at 9DIV (Figure 4E-G). These data show that, in contrast to Np65-GFP, Np65Δ-GFP did neither rescue impaired spinogenesis in *Nptn^-/-^* neurons nor increase the number of dendritic protrusions in rat neurons. Thus, independently of the rodent model from which neurons were derived, Np65 requires its TRAF6 motif to regulate the density of spinogenic protrusions in hippocampal neurons.

**Figure 4.**
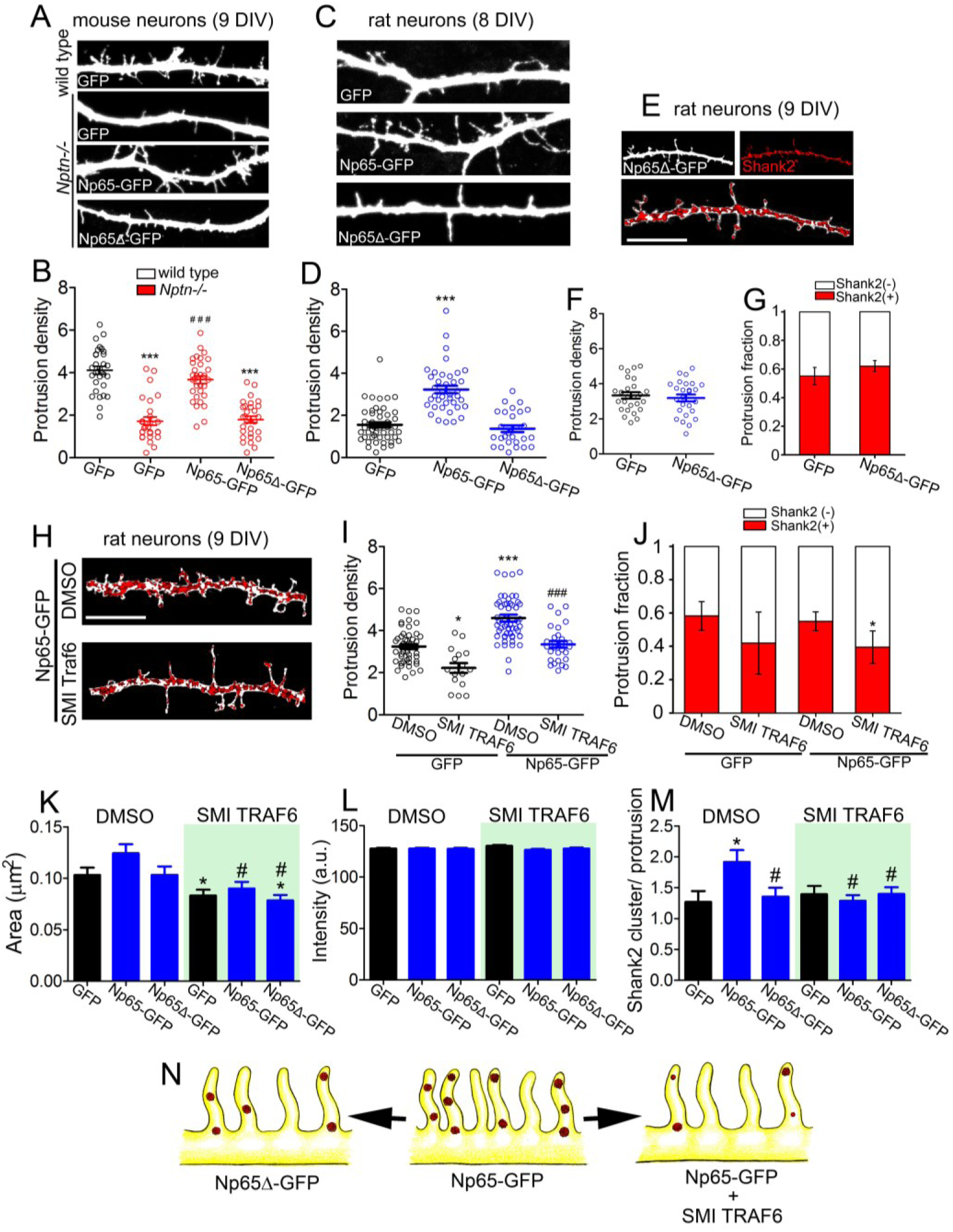
Neuroplastin mediates spinogenesis through TRAF6. **A**, **B**. The TRAF6 binding site-deficient Np65Δ-GFP does not rescue spinogenesis in *Nptn^-/-^* neurons. (**A**) Confocal images of segments of dendrites of *Nptn^+/+^* and *Nptn^-/-^* neurons transfected with plasmids encoding GFP, Np65-GFP or Np65Δ-GFP at 7 DIV. At 9 DIV, these neurons were fixed and stained with anti-GFP antibodies followed by an Alexa 488-conjugated antibody (green). (**B**) Protrusion densities from two independent cultures were used to obtain the mean ± SEM as indicated (*Nptn^+/+^* GFP=4.12 ± 0.18 N=34; *Nptn^-/-^* GFP=1.72 ± 0.19 N=27; *Nptn^-/-^* Np65-GFP=3.67 ± 0.18 N=33; *Nptn^-/-^* Np65Δ-GFP= 1.79 ± 0.16 N=33). ***p<0.001 vs. GFP-filled wild type neurons and ^###^p<0.001 vs. GFP-filled *Nptn^-/-^* neurons using Student‘s t-test. **C**, **D**. Np65Δ-GFP does not foster spinogenesis. (**C**) Dendritic segments of 8 DIV-old rat neurons expressing the indicated proteins upon transfection are shown. (**D**) Protrusion densities from three independent cultures are expressed as the mean ± SEM (GFP=1.72 ± 0.15 N=52, Np65-GFP=3.63 ± 0.11 N=43; Np65Δ-GFP= 1.64 ± 0.16 N=28). ***p<0.001 vs. GFP using Student‘s t-test. **E**-**G**. Expression of Np65Δ-GFP does not promote spinogenesis in 9 DIV-old rat hippocampal neurons. (**E**) Dendritic segments of neurons expressing GFP or Np65Δ-GFP and stained with antibodies against GFP (white) and Shank2 (red clusters) were photographed using confocal microscopy. Scale bar=10 µm. (**F**) Quantification of the protrusion densities and (**G**) the distribution of Shank2-positve and Shank2-negative protrusions from 20-30 neurons per group from three independent cultures (Shank2(+): GFP=0.55 ± 0.06; Np65Δ-GFP=0.62 ± 0.04). **H**-**J**. TRAF6 inhibition decreases spinogenesis. (**H**) 7 DIV-old rat neurons transfected with Np65-GFP and treated with the TRAF6 inhibitor SMI 6860766 (SMI TRAF6, 2 µm) for 48 hours were fixed and stained for GFP (white) and Shank2 (red clusters). Scale bar=10 µm. (**I**) Protrusion density (DMSO GFP=3.24 ± 0.118 N=47; SMI TRAF6 GFP=2.22 ± 0.23 N=16; DMSO Np65-GFP=4.59 ± 0.16 N=56; SMI TRAF6 Np65-GFP=3.34 ± 0.16 N=28) and (**J**) Distribution of Shank2-positve and Shank2-negative protrusions from transfected neurons per group from three independent cultures are displayed. *p<0.05 or ***p<0.001 vs. DMSO GFP and ^###^p<0.001 vs. SMI TRAF6 GFP using Student‘s t-test (Shank2(+): GFP DMSO=0.58 ± 0.08; SMI TRAF6=0.52 ± 0.19; Np65-GFP DMSO=0.55 ± 0.06; Np65-GFP SMI TRAF6=0.42 ± 0.09). **K**-**M**. We calculated (**K**) the area (DMSO GFP=0.105 ± 0.004; DMSO Np65-GFP=0.137 ± 0.004; DMSO Np65Δ-GFP=0.106 ± 0.003; SMI TRAF6 GFP=0.093 ± 0.003; SMI TRAF6 Np65-GFP=0.094 ± 0.006; SMI TRAF6 Np65Δ-GFP=0.098 ± 0.005), (**L**) the fluorescence intensity (DMSO GFP= 134.6 ± 1.4; DMSO Np65-GFP= 139.5 ± 1.9; DMSO Np65Δ-GFP= 138.4 ± 2.1; SMI TRAF6 GFP= 133.0 ± 1.7; SMI TRAF6 Np65-GFP= 134.0 ± 1.8; SMI TRAF6 Np65Δ-GFP= 134.3 ± 1.6), and (**M**) the number of Shank2 clusters per protrusion (DMSO GFP=1.36 ± 0.17; DMSO Np65-GFP=1.92 ± 0.14; DMSO Np65Δ-GFP=1.35 ± 0.14; SMI TRAF6 GFP=1.39 ± 0.13; SMI TRAF6 Np65-GFP=1.28 ± 0.09; SMI TRAF6 Np65Δ-GFP=1.40 ± 0.10) of the experiments in Figure 4H-J. *p<0.05 between Np65-GFP-expressing and GFP-expressing neurons using Student‘s t-test. #p<0.05 between the treatments for the same transfection. **N**. Neuroplastin requires both its TRAF6 binding site and endogenous TRAF6 activity to promote spinogenic protrusion density. The illustration in the middle shows Np65-GFP-expressing neurons with increased density of Shank2-containing spinogenic protrusions. This phenotype is no longer observed when the TRAF6 binding site is deleted from the Np65 intracellular tail (Np65Δ-GFP, left). Incubation with SMI 6860766 (SMI TRAF6) decreases both the density of protrusions and fraction of protrusions with Shank2 clusters (right).

Next, we evaluated whether endogenous TRAF6 is involved in neuroplastin-mediated spinogenesis in rat hippocampal neurons at 9 DIV. We used the small molecule inhibitor (SMI) 6860766, which binds the TRAF-C domain of TRAF6 and blockades its capacity to interact with its binding sites (van den Berg et al., 2014; Chatzigeorgiou et al., 2015) (for simplicity hereafter called SMI TRAF6). Addition of SMI TRAF6 (2 μM) reduced the density of protrusions in GFP- and in Np65-GFP-expressing neurons compared to vehicle-treated (0.01% DMSO) GFP- and Np65-GFP-expressing neurons, respectively (Figure 4H,I). SMI TRAF6 application did not decrease significantly the distribution of Shank2-positive vs. Shank2-negative protrusions in GFP-filled neurons (Figure 4J). However, treatment with SMI TRAF6 decreased the fraction of Shank2-positive protrusions in Np65-GFP-expressing neurons slightly but significantly (Figure 4J). SMI TRAF6 also decreased the area, but not the intensity of Shank2 clusters, and it reduced the number of Shank2 clusters per protrusion in Np65-GFP-expressing neurons to the level of controls (Figure 4K-M). Moreover, SMI TRAF6 treatment evidenced that the size of Shank2 clusters depends on TRAF6 (Figure 4K). We conclude that neuroplastin strictly requires both its TRAF6 binding site and TRAF6 to increase the density of dendritic protrusions in hippocampal neurons. Either deficiency of these pre-requisites abrogates the spinogenic capacity of neuroplastin (Figure 4N).

Very recently, it has been discovered that neuroplastin interacts with all four plasma membrane Ca^2+^ ATPases (PMCA1-4) stabilizing the surface expression of these pumps (Bhattacharya et al., 2017; Herrera-Molina et al., 2017; Korthals et al., 2017; Schmidt et al., 2017; Gong et al., 2018). Thus, we addressed the obvious question whether TRAF6 or neuroplastin require PMCA to promote dendritic protrusion density. First, we examined whether the TRAF6 binding motif of neuroplastin is necessary to maintain PMCA protein. Consistent with our previous report (Herrera-Molina et al., 2017), full-length Np65-GFP increased protein levels of PMCA2 compared to GFP when co-transfected in HEK cells (Figure S5A,B). Np65Δ-GFP was found as effective as Np65-GFP to increase PMCA2 levels in co-transfected HEK cells (Figure S5A,B). In rat hippocampal neurons at 9 DIV, confocal microscopy revealed that both Np65-GFP and Np65Δ-GFP co-localized with endogenous PMCA and increased PMCA protein levels (Figure S5C,D). Also, SMI TRAF6 treatment from 7-9 DIV did not modified total PMCA protein levels in young hippocampal neurons (data not shown). Therefore, either elimination of the TRAF6 binding motif of neuroplastin or SMI TRAF6 treatment abrogates TRAF6-mediated spinogenetic function (Figure 4), but does not affect endogenous PMCA protein levels or the capacity of neuroplastin to promote the expression levels of the pump. Although PMCA inhibition with Caloxin 2a1 seemed to be slightly enlarge protrusions, the density of protrusions was not affected in GFP-filled (Figure S5E,F) nor in Np65-GFP-expressing neurons at 9 DIV (data not shown). Thus, PMCA is not critically required by TRAF6 to mediate neuroplastin-promoted spinogenesis.

### Blockage of TRAF6 impairs formation of excitatory synapses and impacts their activity

Although, TRAF6 seems to be essential for brain development (Lomaga et al., 2000) and plasticity of mature synapses (Ma et al., 2017), it has never being linked to synapse formation. Based our observation that TRAF6 mediates spinogenesis by neuroplastin; we wondered whether direct blockage of TRAF6 with SMI 6860766 during spinogenesis in young neurons alters the number of formed excitatory synapses when neurons mature. To test this, rat hippocampal neurons were treated with SMI TRAF6 (2 μM) or with vehicle (0.01% DMSO) from 7 to 9 DIV and analyzed at 12 DIV (Figure 5A). The number of excitatory synapses (homer-positive spines matching synapsin-positive presynapses, Herrera-Molina et al., 2014) per 10 µm dendrite was significantly reduced in neurons treated with SMI TRAF6 from 7 to 9 DIV (Figure 5B) as compared to vehicle-treated neurons. In contrast, neurons treated with SMI TRAF6 from 10 to 12 DIV and analyzed at 12 DIV displayed similar number of synapses as vehicle-treated neurons (Figure 5B). These data indicate that TRAF6 is critical for the early spinogenesis of some ∼25% of hippocampal excitatory synapses *in vitro*.

**Figure 5.**
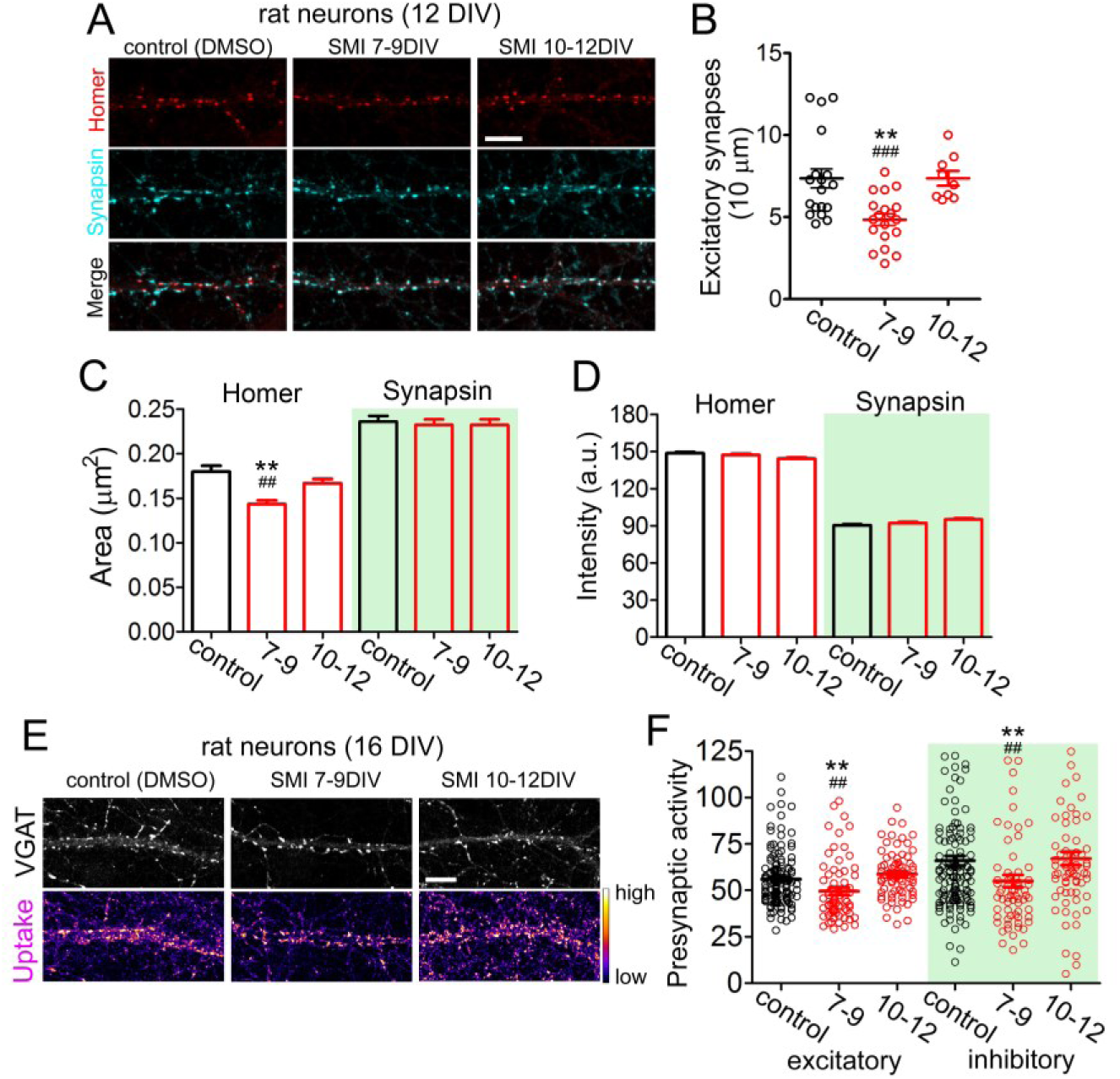
TRAF6 blockage during early, but not late, spinogenesis decreases excitatory synapse density and impairs balanced E/I ratio. **A**-**D**. Treatment with SMI 6860766 (SMI TRAF6) from 7 to 9 DIV, but not from 10 to 12 DIV reduces the number of excitatory synapses. (**A**) Representative confocal images of dendritic segments stained with antibodies against synaptic markers (red, postsynaptic Homer; cyan, presynaptic Synapsin-1) at 12 DIV. As indicated, rat hippocampal neurons were previously treated with SMI TRAF6 for 48 hours between days 7 to 9 or 10 to 12. Scale bar=10 µm (**B**) Quantification of the number excitatory synapses per 10 µm of dendritic segment from two independent experiments (control=7.36 ± 0.58 N=19; 7-9=4.84 ± 0.35 N=19; 10-12=7.36 ± 0.45 N=9). **p<0.01 vs. control and ^###^p<0.001 vs. 10-12 using Student‘s t-test. (**C**) Quantification of the area (Homer: control=0.172 ± 0.004; 7-9=0.143 ± 0.004; 10-12=0.162 ± 0.006. Synapsin: control=0.236 ± 0.006; 7-9=0.232 ± 0.007; 10-12=0.248 ± 0.003) and (**D**) fluorescence intensity (Homer: control=148.58 ± 0.99; 7-9=147.13 ± 0.99; 10-12=145.58 ± 1.12. Synapsin: control=90.37 ± 0.79; 7-9=92.21 ± 0.84; 10-12=96.77 ± 2.01) of homer- and synapsin-positive puncta. For Homer, **p<0.01 vs. control and ^##^p<0.01 vs. 10-12 using Student‘s t-test. **E**, **F**. Reduced presynaptic vesicle recycling in SMI 6860766 (SMI TRAF6)-treated 16 DIV-old neurons. (**E**) Dendritic segments of rat neurons, treated with SMI TRAF6 when indicated, tested for uptake of fluorescently labelled antibody against the luminal part of the presynaptic protein synaptotagmin at 16 DIV (see Material and Methods), fixed, stained with an anti-VGAT antibody to discriminate inhibitory presynapses from the excitatory ones. Photomicrographs with a confocal microscope are shown. The heat scale bar indicates uptake levels. Scale bar=10 µm. (**F**) Quantification of presynaptic activities from three independent experiments (Excitatory: control=55.97 ± 1.28 N=130; 7-9=49.58 ± 2.025 N=67; 10-12=58.73 ± 1.568 N=69; Inhibitory: control=66.00 ± 2.63 N=130; 7-9=54.99 ± 3.30 N=67, 10-12=67.11 ± 3.62 N=69). **p<0.01 vs. control and ^##^p<0.01 between synapse type using Student‘s t-test.

Next, we quantified the area and fluorescence intensity of the homer-positive and synapsin-positive puncta that were still present after the treatment with SMI TRAF6. The area of homer-positive puncta was reduced in rat neurons treated with SMI TRAF6 compared to control vehicle-treated rat neurons from 7 to 9 DIV. Neurons treated with SMI TRAF6 from 10 to 12 DIV displayed similar area of homer clusters compared to control vehicle-treated neurons (Figure 5C). The intensity of homer-positive puncta as well as the area and intensity of synapsin-positive puncta were not affected by SMI TRAF6 (Figure 5C-D). Thus, TRAF6 inhibition during spinogenesis from 7 to 9 DIV decreases the enrichment of the postsynaptic marker homer, but not of the presynaptic protein synapsin. This phenotype resulting from TRAF6 inhibition is very similar to the one previously reported in neuroplastin-deficient neurons (Herrera-Molina et al., 2014).

TRAF6 is critical for early spinogenesis of Shank2-containing spines at 9 DIV (Figure 4 and Figure S3) and for the formation of excitatory synapses at 12 DIV (Figure 5). Thus, we tested whether the presynaptic activity is consequentially affected by TRAF6 inhibition in mature synapses. Not surprisingly, evaluation of the presynaptic uptake of synaptotagmin-1 antibody – reporting vesicle release and recycling driven by intrinsic network activity – showed a decreased activity in mature VGAT-negative excitatory presynapses after treatment with SMI TRAF6 from 7 to 9 DIV, but not from 10 to 12 DIV (Figure 5F). Therefore, in addition to the anatomical evidence (Figure 4, Figure S4, and Figure 5), this result shows that spinogenetic function of TRAF6 in young neurons is critically linked to a long-term establishment of a balanced E/ I synapse activity in mature neurons.

Interestingly, the activity of VGAT-positive inhibitory terminals was also reduced in at 16 DIV when neurons were treated with SMI TRAF6 from 7 to 9 DIV, but not from 10 to 12 DIV (Figure 5F). Similarly, VGAT-positive inhibitory terminals placed on the soma of neurons displayed decreased activity after the treatment with SMI TRAF6 from 7 to 9 DIV, but not from 10 to 12 DIV (not shown). To unravel the nature of these results, we calculated the area of vesicle release (mean area of puncta) and the activity level (mean intensity per pixel) for each presynapse type. From these data (Table 2), we deduced that decreased inhibitory activity results from an adaptation to the weakening of the glutamatergic activity due to the reduction of the density of spinogenic protrusions by earlier TRAF6 blockage (Figures 1, 4, S4, and 5). Consistent with this explanation, TRAF6 inhibition after spinogenesis did not affect the number of excitatory synapses (Figure 5).

**Table 2.**
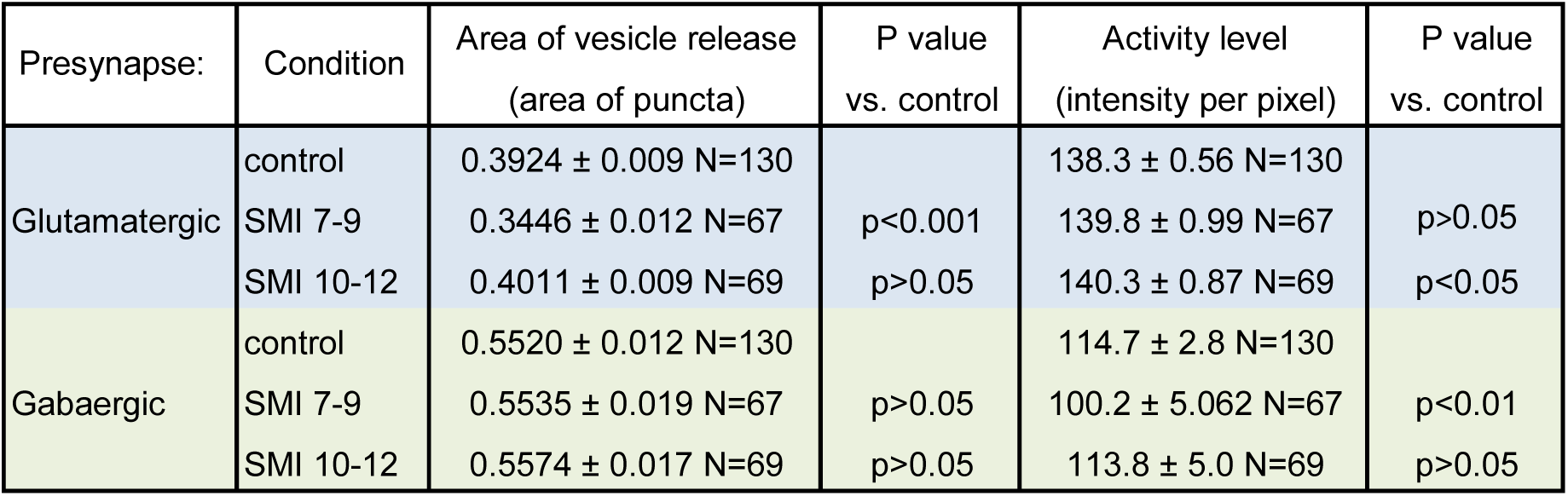
Synaptotagmin-1 assay in inhibitory and excitatory presynapses. The table is explained in the main text and in Figure 5.

| Presynapse: | Condition | Area of vesicle release<br>(area of puncta) | P value<br>vs. control | Activity level<br>(intensity per pixel) | P value<br>vs. control |
| --- | --- | --- | --- | --- | --- |
| Glutamatergic | control | 0.3924 ± 0.009 N=130 |  | 138.3 ± 0.56 N=130 |  |
|  | SMI 7-9 | 0.3446 ± 0.012 N=67 | p<0.001 | 139.8 ± 0.99 N=67 | p>0.05 |
|  | SMI 10-12 | 0.4011 ± 0.009 N=69 | p>0.05 | 140.3 ± 0.87 N=69 | p<0.05 |
| Gabaergic | control | 0.5520 ± 0.012 N=130 |  | 114.7 ± 2.8 N=130 |  |
|  | SMI 7-9 | 0.5535 ± 0.019 N=67 | p>0.05 | 100.2 ± 5.062 N=67 | p<0.01 |
|  | SMI 10-12 | 0.5574 ± 0.017 N=69 | p>0.05 | 113.8 ± 5.0 N=69 | p>0.05 |

### TRAF6 is required for synapse transmission and network-driven neuronal activity

We evaluated whether the defined function of TRAF6 in young neurons undergoing early spinogenesis has a long standing impact on synapse transmission when neurons have matured. Hippocampal neurons were treated SMI TRAF6 or vehicle from 6 to 9 DIV, leaved to mature, and impaled to record intracellularly miniature excitatory postsynaptic currents (mEPSCs) using Patch Clamp technique in the presence of 1µM TTX at 18-23 DIV (Figure 6A). In SMI TRAF6-treated neurons, both amplitude and decay time of mEPSCs were altered whereas rise time remained unchanged compared to vehicle-treated neurons (Figure 6B) indicating physiological alterations at the postsynaptic levels. These experiments are in tight agreement with those describing the time period of the TRAF6-Np-dependent-spinogenesis (Table 1; Figs. 1 and 4) as well as they are coherent with the anatomical results showing that TRAF6-controlled spinogenesis *via* Np in young neurons is require for the long-term establishment of a proper synapse number as neurons mature (Figure 5).

**Figure 6.**
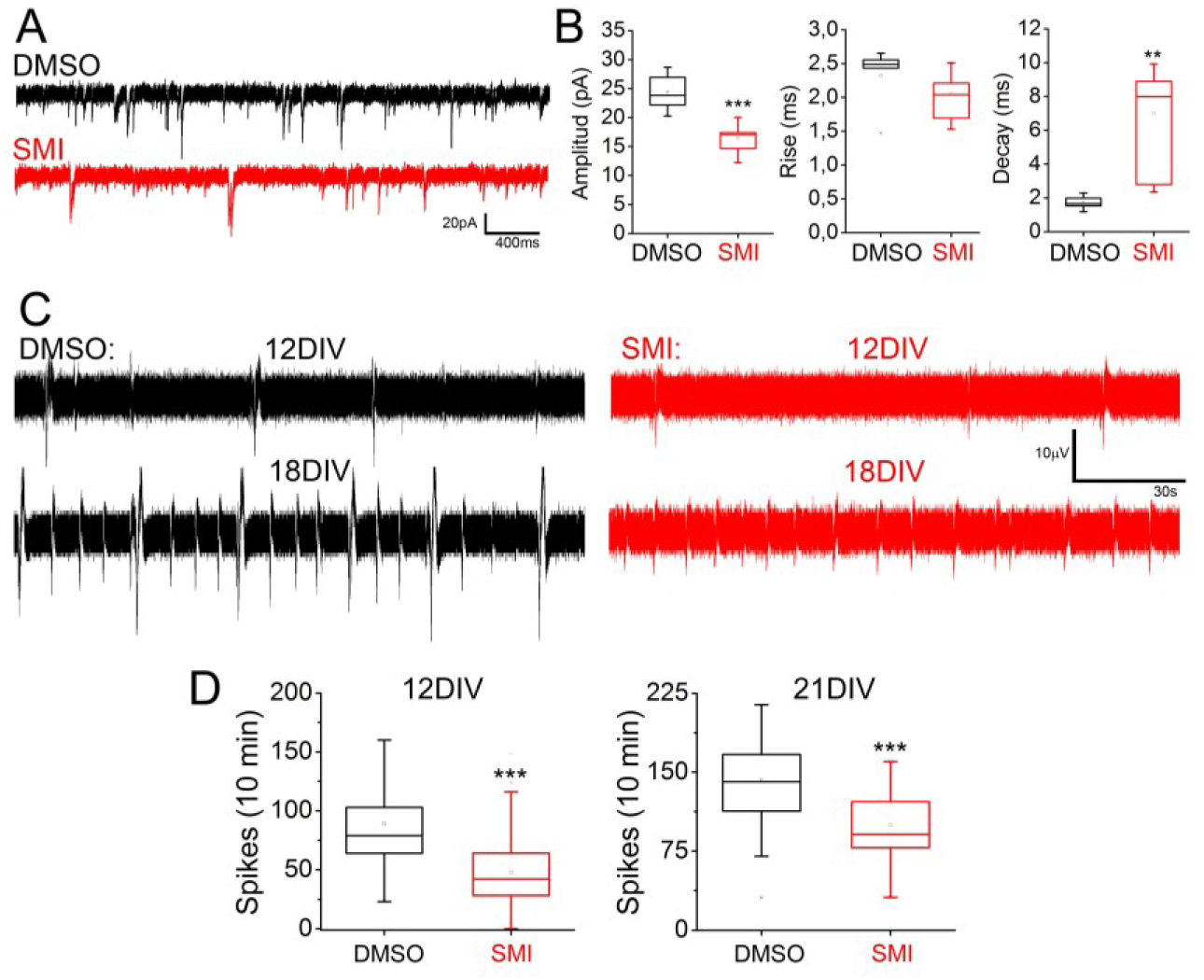
TRAF6 function is necessary for electrophysiological maturation of neurons. **A**, **B**. Treatment with SMI TRAF6 from 6 to 9 DIV impairs mEPCSs in 18-23 DIV hippocampal neurons. (**A**) Representative traces of intracellular recordings of mEPCSs. (**B**) Quantification of the amplitude (DMSO=24.389 ± 1.531; SMI=16.384 ± 0.829), rise time (DMSO=2.320 ± 0.2150; SMI=2.054 ± 0,113),, and decay time (DMSO=1.729 ± 0.190; SMI=7.008 ± 1.039) of mEPCSs of 12 DMSO-treated and 14 SMI-treated neurons from 4 independent cultures. ***p<0.001 or ** p<0.01 vs. DMSO using Student‘s t-test. **C**-**D**. Reduced activity in hippocampal neurons treated with SMI TRAF6 from 6 to 9 DIV. (**C**) Traces of extracellularly-recorded neuronal activity obtained consecutively when neurons were 12 and also 18 DIV. (**D**) Quantification of the number of spikes per electrode were obtained using Matlab (12DIV: DMSO= 89,2 ± 3,5, SMI= 47,6 ± 1,9; 18DIV: DMSO= 142,5 ± 2,8, SMI= 79,5 ± 2,2).

To confirm further the functional relevance of our findings for neuronal physiology, we evaluated the effect of TRAF6 blockage with SMI TRAF6 from 6 to 9 DIV on network-driven activity of hippocampal neurons cultured in multi-electrode arrays at 12 and 18 DIV (Figure 6C). As expected from the anatomical and electrophysiological evidence (see before), neurons in SMI TRAF6-treated arrays displayed lower number of extracellular spikes compared to neurons in control arrays at 12 and 18 DIV (Figure 6D). Therefore, blockage of TRAF6 in early spinogenesis has a long-lasting and permanent impact on neuronal activity.

## Discussion

Our study addresses the question of how neurons develop early the capacity to form an adequate number of well-functioning synapses to communicate with each other. We show that TRAF6 controls glutamatergic synaptogenesis critically relevant for the correct establishment of excitatory synapse density instructing E/I synapse balance, synaptic transmission, and neuronal activity. We discover that TRAF6 function in early spinogenesis is directly linked to its interaction with neuroplastin which has been reported to be somehow related to synapse formation *in vivo* (Herrera-Molina et al. 2014; Amuti et al., 2016; Carrott et al., 2016; Zeng et al., 2016; Bhattacharya et al., 2017).

We found TRAF6 to be necessary for early excitatory spinogenesis revealing a novel function for this factor in neuronal development. This function is different from the one related to failed p75 neurotrophin receptors (p75NTR)-regulated apoptosis and developmental alterations in the morphogenesis of TRAF6 KO embryonal brains (Lomaga et al., 2000; Yeiser et al., 2004). Indeed, our data demonstrate that TRAF6 is effective to mediate neuroplastin-dependent spinogenesis during a defined time window in neuronal development i.e. when most dendritic protrusions are forming and when regularly spaced and cell membrane-associated co-localization of TRAF6 with neuroplastin in dendritic protrusions occurs. Later in mature neurons, TRAF6 did not co-localize with neuroplastin and neither TRAF6 inhibition nor neuroplastin overexpression modified the density of protrusions or the number of excitatory synapses. Instead, it has been reported recently that TRAF6 efficiently binds to and regulates the stability of PSD-95 as a necessary step for the synaptic plasticity of mature neurons (Ma et al., 2017). This can explain the developmental shift of the binding of TRAF6 with neuroplastin in young neurons towards PSD-95 a much more prominent interaction partner in mature neurons. Accordantly, PSD-95 levels are lower than neuroplastin levels in young neurons (Buckby et al., 2004) but highly abundant in glutamatergic spines of mature neurons. Additionally, neuroplastin could also change its predilection for interaction partners in young neurons as it becomes an essential subunit for the four paralogs of PMCA in mature neurons (Bhattacharya et al., 2017; Herrera-Molina et al., 2017; Korthals et al., 2017; Schmidt et al., 2017; Gong et al., 2018). Therefore, our report comes to fill the gap between the functions of TRAF6 in early neuronal survival and synaptic plasticity in mature neurons.

We demonstrated here that neuroplastin harbor a single intracellular TRAF6 binding site. The intracellular sequence RKRPDEVPD of neuroplastin fulfilled structural and three-dimensional criteria to be considered as a proper TRAF6 binding site (Ye et al., 2002; Sorrentino et al., 2008; Yin et al., 2009). The binding affinity between neuroplastin-derived peptide carrying the TRAF6 binding site with TRAF6 C-terminal domain was very close to the expected one according to solid crystallographic, structural and functional data demonstrating the specialization of the TRAF6 binding site for binding TRAF6 (Ye et al., 2002; Sorrentino et al., 2008; Yin et al., 2009). Co-precipitation between the neuroplastin isoforms with TRAF6 was not affected by the presence of the alternative splicing DDEP proximal to the TRAF6 binding site. However, it was drastically reduced or eliminated by mutation or deletion of the TRAF6 binding site of neuroplastin. Endogenous TRAF6 and GFP-tagged TRAF6 were recruited into the regularly spaced cell membrane-associated puncta by neuroplastin only when an intact TRAF6 binding site was present. This is coherent with TRAF6 moving towards the cell membrane and forming micrometric and geometrically organized lattice-like supramolecular structures able to regulate the multimerization of bound transmembrane proteins and to host downstream cell signaling elements (Schultheiss et al., 2001; Yin et al., 2009; Ferrao et al., 2012; Wu et al., 2013). Elimination of the lattice-forming RING domain did not prevented TRAF6 recruitment by neuroplastin but abrogated the capacity of the transmembrane glycoprotein to promote the formation of filopodial structures. Thus, it is realistic to propose that upon TRAF6 binding and higher-order oligomerization of the factor, neuroplastin might become a part of such supramolecular complexes to initiate downstream events of cell signaling causing the formation of filopodial structures in HEK cells and spinogenic protrusion in neurons.

The selective and specific binding of TRAF6 places neuroplastin as novel candidate to organize prolific signaling mechanisms related to spinogenesis and/or to stabilization young spines. Indeed, preliminary experiments suggest that inhibition of p38 MAPK, ERK1/2 or PI3 kinase reduces the number of neuroplastin-promoted filopodia in HEK cells and dendritic protrusions in young hippocampal neurons (Vemula and Herrera-Molina, unpublished data). It has been reported that extracellular engagement of neuroplastin activates p38 MAPK (Empson et al., 2006), ERK1/2 and PI3 kinase (Owczarek et al., 2011). These signalling pathways could also be related to homophilic *trans*-synaptic engagement of Np65 to foster and or stabilize young protrusion. The literature recognized TRAF6 largely as a main upstream activator of transcriptional factor NFκB pathway (Darnay et al., 1999; Xie 2013). In young hippocampal neurons, NFκB activity is not regulated by neuronal activity but, it is necessary for the formation of excitatory synapses (Boersma et al., 2011). Also, the constitutively high NFκB activity in young neurons maintains glutamatergic spinogenesis contributing in turn to the future establishment of E/I synapse balance in mature neurons (Boersma et al., 2011; Dresselhaus et al., 2018). Preliminary experiments suggest that inhibition of NFκB nuclear translocation reduces the density of neuroplastin-induced dendritic protrusions in young hippocampal neurons. Thus, it is possible that neuroplastin regulates NFκB activity in a TRAF6-dependent manner as a necessary step to promote the formation of filopodial structures in HEK cells and spinogenic protrusions in hippocampal neurons – a hypothesis that needs to be tested in future.

TRAF6 is strictly required by both isoforms of neuroplastin to promote spinogenesis in young hippocampal neurons. Expression of Np55 or Np65 equally rescued the reduced number of dendritic protrusions in *Nptn^-/-^* neurons and enhanced normal formation of spinogenic protrusions in rat neurons upon overexpression. This implies that TRAF6 is a spinogenic mechanism that do not essentially dependent of the Np65-specific *trans*-adhesive extracellular domain (Smalla et al., 2000). In mouse models, elimination of both Np55 and Np65 is required to reduce the density of hippocampal excitatory synapses (Herrera-Molina et al., 2014, Amuti et al., 2016; Bhattacharya et al., 2017) and ribbon synapses (Carrott et al., 2016). In contrast, elimination of Np65 only is not sufficient to alter the synapse density; however, it results in morphological alterations of hippocampal spines in Np65 KO mice (Amuti et al., 2016). These data, however, do not rule out the possibility of a later participation of the Np65-specific adhesive Ig-like domain in the *trans*-stabilization of postsynapses (Smalla et al., 2000; Herrera-Molina et al., 2014). This needs to be addressed by future experiments. The present finding that over-expression of Np65 fosters the formation Shank2-containing protrusions indicates that these additional and newly formed structures are spinogenic in transfected neurons under development. Additionally, this finding matches with previous observations, i.e. a tight correlation between the fast up-regulation of expression of neuroplastin and intracellular synaptic proteins during early synaptogenesis in cultured and in acutely isolated rat hippocampal slices (Buckby et al., 2004) as well as the stabilization and maturation of spines (Böckers et al., 1999; Sarowar and Grabrucker 2016).

Because TRAF6 binding motif in not present in most spinogenic CAMs, TRAF6 mediated spinogenesis seems to be rather restricted to neuroplastin. Cell adhesion molecules have been proposed as key participants in synapse formation. Nonetheless, knocking out *trans-*synaptic CAMs, that are crucially involved in synapse maturation, transmission and/or plasticity, does not significantly affect the number of synapses formed *in vivo* (Henderson et al., 2001; Missler et al.; 2003; Varoqueaux et al, 2006; Chubykin et al., 2007; Linhoff et al., 2009; Bozdagi et al., 2010; Robbins et al, 2010). Thus, it has been considered that still unexplored molecular machineries interplaying with CAMs may promote synapse formation (Yoshihara et al., 2009; Jiang et al., 2017; Jang et al., 2017; Südhof 2017). Independent of their adhesive properties some CAMs can form transmembrane complexes in *cis* that in turn can recruit intracellular proteins and activate signaling mechanisms underlying neuronal development (Cavallaro and Dejana, 2011; Jang et al., 2017). Our results shown that neuroplastin contributes to the formation of spinogenic dendritic protrusions cell-autonomously and, the binding of TRAF6 confers to neuroplastin unique mechanistic possibilities to potentially regulate cell signalling and gene expression related to control spinogenesis.

## Materials and Methods

### Cells

Primary *Nptn^-/-^* neurons were derived from hippocampi of *Nptn^-/-^* mice and compared to primary *Nptn^+/+^ neurons* derived from their proper control *Nptn^+/+^* mice (Herrera-Molina et al., 2014; Bhattacharya et al., 2017). Cultures of rat hippocampal neurons were obtained as described (Herrera-Molina et al., 2005; 2012). Human embryonic kidney (HEK) 293T cells were cultured as previously described (Herrera-Molina et al., 2017).

### DNA constructs and transfections

GFP-tagged neuroplastin constructs have been described (Herrera-Molina et al., 2017). Neuroplastin mutants flanked by HindIII and BamH1 restriction sites were generated from Np65-GFP plasmid by PCR amplification using the following primers for Np65-GFP forward: 5’-TCA AGC TTG CCA CCA TGT CG-3’ reverse: 5’-GGC GAT GGA TCC ATT TGT GTT TC-3’; Np65Δ-GFP reverse 5’-GGA TCC TGG CCT CTT CCT CTT CTC ATA C-3’: Np65_PED_-GFP forward 5’-GAG GAA GAG GGC AGA TGC GGT TCC TGC TG-3’ reverse 5’-CAG CAG GAA CCG CAT CTG CCC TCT TCC TC-3’. The mouse N-terminal Flag-tagged TRAF6 (Flag-TRAF6) mammalian expression plasmid was purchased from Addgene (#21624, GenBank: BAA12705.1). N-terminally GST-tagged TRAF6 (GST-TRAF6) and RING domain deficient TRAF6 with coiled-coil domain and TRAF6-C domain (289-530aa; GST-TRAF6_cc-c_) plasmids with BamH1 and EcoR1 restriction sites were generated by PCR amplification. GST-TRAF6 forward 5’-GAC AGG ATC CTC ATG AGT CTC TTA AAC-3’ reverse 5-TAC GAA TTC CTA CAC CCC CGC ATC AGT A-3’; GST-TRAF6_cc-c_ forward 5-GCG TCG GAT CCA TAT GGC CGC CTC T-3’; TRAF6-GFP forward 5’-GTG AAG CTTCTA ATG AGT CTC TTA AAC TGT GA-3’ reverse 5’-ATA AGG ATC CCT ACA CCC CCG CAT C-3’; TRAF6_cc-c_-GFP forward 5’-GTG AAG CTT CTA ATG GCC GCC TCT-3’. Scrambled siRNA (sc-37007) and TRAF6 siRNA (sc-36717) were purchased from Santa Cruz. HEK cells were transiently transfected with plasmid DNA constructs using Lipofectamine 2000 (Invitrogen/ThermoFisher) in optiMEM media (Gibco). Primary neuronal cultures were transfected at 7 days *in vitro* (DIV). Control scrambled siRNA or TRAF6 siRNA (30 nM) were transfected using siLentFect (Bio-Rad).

### *In silico* modelling

We performed local peptide docking based on interaction similarity and energy optimization as implemented in the GalaxyPepDock docking tool (Lee et al., 2015). The protein–peptide complex structure of the hTRANCE-R peptide bound to the TRAF6 protein as provided by Ye et al., 2002 was used as input (PDB: 1LB5). The docking employs constraints of local regions of the TRAF6 surface based on the interaction template. The energy-based optimization algorithm of the docking tool allows efficient sampling of the backbone and side-chains in the conformational space thus dealing with the structural differences between the template and target complexes. Models were sorted according to protein structure similarity, interaction similarity, and estimated accuracy. The fraction of correctly predicted binding site residues and the template-target similarity was used in a linear model to estimate the prediction accuracy. The model using target-template interactions based on the QMPTEDEY motif of the hTRANCE-R template was selected (TM score: 0.991; Interaction similarity score 108.0; Estimated accuracy: 0.868).

### Surface plasmon resonance

Protein-Protein interaction measurements were carried out on a BIACORE X100 (GE Healthcare Life Sciences). Sensorgrams were obtained as single cycle kinetics runs. Therefore increasing concentrations of neuroplastin peptide (2.5, 5, 100, 200, 400µM) or just running buffer (startup) were sequentially injected on GST-TRAF6 coated CM5 sensor chip (GE). Unspecific bindings were calculated by using a GST-coated sensor as reference response. Immobilization of these proteins was done using the amine coupling kit as we described in Reddy et al. 2014. All runs were performed in HBS-P buffer. Analysis of affinity was performed using the BIACORE X100 Evaluation Software 2.0.1 (Reddy et al. 2014).

### GST pull-down assay

GST, GST-TRAF6 and GST-TRAF6_cc-c_ were transformed into *Escherichia coli* BL21 (DE3) bacterial strain and induced by 0.5 mM of isopropyl-1-thio-b-D-galactopyranoside (IPTG) for 6 h at 25°C. The cells were lysed in resuspension buffer (50 mM Tris-HCl pH 8.0, 150 mM NaCl and protease inhibitor cocktail (Roche) with sonication on ice. The purifications of these proteins from transformed bacterial cell extract were performed according to manufacturer instructions (GST bulk kit, GE Healthcare Life Sciences). The purified soluble GST proteins were immobilized on glutathione sepharose 4B beads (GE Healthcare Life Sciences). The beads were washed with binding buffer at least four times, and the pull-down samples were subsequently subjected to immunoblot analyses. The 5 μg of fusion protein coupled beads (GST, GST-TRAF6 and GST-TRAF6_cc-c_) were incubated with lysate from HEK cells transfected with Np65-GFP for 1 h at 4°C in 500 μl RIPA lysis buffer. The beads were washed and eluted with pre-warmed SDS sample buffer. The eluted complexes were resolved by SDS-PAGE.

### Immunoblotting

Proteins were separated by sodium dodecyl sulfate– polyacrylamide gel electrophoresis (SDS-PAGE) on 10% gels and transferred to a nitrocellulose membrane (Whatman). After blocking with 5% non-fat milk in Tris-buffered saline (TBS) containing 0.1% of Tween 20 for 1 h, the membranes were incubated with indicated antibodies overnight, washed with TBS three times, and then incubated with corresponding secondary antibody conjugated to horseradish peroxidase enzyme for 1 h. Immunodetection was performed with the following antibodies: anti-Flag mouse (Sigma, #F1804; 1:2,000), anti-GFP rabbit (Abcam, #ab290; 1:2,500), anti-TRAF6 mouse (Santa Cruz, #sc-8709; 1:1,000) and anti-β-actin mouse (Sigma, #A5441; 1:1,000), horseradish peroxidase-conjugated anti-mouse (Dako, #P0447; 1:4,000) or anti-rabbit IgG (gamma-chain specific, Sigma, #A1949-1VL; 1:4,000) antibodies.

### Immunocytochemistry

Hippocampal neurons were fixed with 4% PFA for 8 mins and then washed with a solution containing 10% horse serum, 0.1 mM glycine, and 0.1% Triton X-100 in Hanks’ balanced salt solution four times for 5 min. Fixed samples were incubated with indicated primary antibodies for overnight at 4°C. To visualize dendritic protrusions, after transfection, pyramidal neurons were morphologically identified based on the side and shape of cell body as observed using anti-MAP2 guinea pig (Synaptic Systems, #188 004; 1:1,000) and anti-GFP mouse (Sigma Aldrich, #11814460001; 1:1,000) antibodies. Routinely, neuron identity was confirmed using an anti-Ctip2 rat (Abcam, #25B6; 1:250) (Herrera-Molina et al., 2014). Subsequently, samples were incubated with anti-guinea pig Cy5-, anti-mouse Alexa 488-conjugated secondary antibodies (1:1,000) generated in donkey (Jackson ImmunoResearch) for 1 hour at RT. Other primary antibodies used were: anti-Synapsin 1 rabbit (Synaptic Systems, #106 103; 1:500), anti-Shank2 guinea pig antibody (Synaptic Systems, #162 204; 1:1,000), anti-Homer1 mouse (Synaptic Systems, #160 011; 1:500), anti-RelA (p65) rabbit (Santa Cruz, #sc-372; 1:500); anti-MAP2 guinea pig (Synaptic Systems, #188 004; 1:1,000) primary antibodies for overnight at 4°C. Subsequently, samples were incubated with anti-rabbit 405-, anti-mouse Cy5-, anti-rat Alexa 488- and/ or anti-guinea pig Cy3-conjugated donkey secondary antibodies (1:1,000) for 1 h. Then samples were washed and mounted with Mowiol. Quantification of synapse marker signals was performed as in detail described in Herrera-Molina et al., 2014.

### Image acquisition and quantification of filopodia/ dendritic protrusions

Images were acquired using HCX APO 63/1.40 NA or 100/1.4NA objectives coupled to a TCS SP5 confocal microscope under sequential scanning mode with 4.0- to 6.0-fold digital magnification. Z-stacks (41.01 x 41.01 x 5 μm physical lengths) were digitalized in a 512 x 512 pixels format file. In HEK cells, filopodia number and length were quantified using a MATLAB-based algorithm, FiloDetect, with some modifications (Nilufar et al., 2013). The algorithm was run for every single image, and the image threshold was adjusted to avoid false filopodia detection and to quantify precise filopodia length and number. The filopodia number per μm was calculated from perimeter of the cell using ImageJ. In neurons, the dendritic protrusions were quantified manually using maximum intensity and Z-projection method of ImageJ software. The dendritic protrusions were considered between 0.25 μm and 20 μm length. Shank clusters were quantified from cropped images using original GFP fluorescent as reference to identify puncta of interest.

### Co-immunoprecipitation assay

HEK cells overexpressing GFP-tagged constructs were washed in ice-cold PBS and lysed using radioimmunoprecipitation assay (RIPA) buffer contained 20 mM of Tris (pH 7.5), 100 mM of NaCl, 1 mM EDTA, 10% glycerin, 0.1% SDS, 1% Triton X-100, 1 mM AEBSF, 1 mM sodium orthovanadate, 1 mM sodium molybdate, 1 mM N-Ethylmalemide, 20 mM sodium fluoride, 20 mM glycerol-2-phosphate, 10 mM potassium hydrogen phosphate, 10 mM sodium pyrophosphate and protease inhibitor cocktail (Roche). Samples were incubated with GFP antibody-coupled magnetic beads (μMACS) at 4° C for 4 h. Immunoprecipitated complexes were eluted using μMACS GFP isolation kit (#130-091-125) according to manufacturer’s instructions. Eluted complexes were subjected SDS-PAGE.

### Synaptotagmin Uptake Assay

Presynaptic activity driven by endogenous network activity was monitored as described before (Herrera-Molina et al., 2014). Hippocampal neurons were washed once with pre-warmed Tyrodes buffer (119 mM NaCl, 2.5 mM KCl, 25 mM HEPES, pH 7.4, 30 mM glucose, 2 mM MgCl_2_, 2 mM CaCl_2_) and immediately incubated with an Oyster 550-labeled anti-synaptotagmin-1 rabbit antibody (Synaptic Systems, #105 103C3; 1:500) for 20 min at 37°C. After the antibody uptake, neurons were washed, fixed, and stained with anti-VGAT guinea pig (Synaptic Systems, #131 004; 1:1,000) and anti-synaptophysin mouse (company, catalog number; 1:1,100) primary antibodies overnight at 4°C. Subsequently, samples were incubated with anti-rabbit Cy3-, anti-guinea pig Cy5- and anti-mouse Alexa 488-conjugated donkey secondary antibodies (1:1,000) for 1 hour. Z-stack images of soma and secondary/tertiary dendrites were acquired using an oil-immersion (HCX APO 63/1.40 NA) objective coupled to a TCS SP5 confocal microscope under sequential scanning mode with a 4.0-fold digital magnification, and digitalized in a 512 x 512 pixels format file (61.51 x 61.51 μm physical lengths). All parameters were rigorously maintained during the image acquisition. For quantification, z-stacks were projected using “sum slices” Z-projection method of Fiji software. We quantified the synaptotagmin-associated fluorescence co-localizing with 1-bit masks derived from VGAT-positive (inhibitory presynapses) or VGAT-negative synaptophysin-positive (excitatory presynapses) puncta using the “image calculator” in the Fiji software. During image processing the original settings of the synaptotagmin channel were carefully maintained as the original. One-bit masks were generated using the analyze particle in the Fiji software for a segmented image of each presynaptic marker (range of particle size 0.15 – 2.25 μm^2^ for inhibitory presynapses and 0.15 – 1.50 μm^2^ for excitatory presynapses).

### Electrophysiology

Whole-cell patch clamp recordings were performed under visual control using phase contrast and sCMOS camera (PCO panda 4.2). Borosilicate glass pipettes (Sutter Instrument BF100-58-10) with resistances ranging from 3–7 MΩ were pulled using a laser micropipette puller (Sutter Instrument Model P-2000). Electrophysiological recordings from neurons were obtained in Tyrode’s medium ([mM] 150 NaCl, 4 KCl, 2 MgCl_2_, 2 MgCl_2_, 10 D-glucose, 10 HEPES; 320 mOsm; pH adjusted to 7.35 with NaOH and Osmolarity of 320 mOsm) + 0.5 µM TTX (Toris).

Pipettes were filled using standard intracellular solution ([mM] 135 K-gluconate, 4 KCl, 2 NaCl, 10 HEPES, 4 EGTA, 4 MgATP, 0.3 NaGTP; 280 mOsm; pH adjusted to 7.3 with KOH). Whole-cell configuration was confirmed via increase of cell capacitance. During voltage clamp experiments neurons were clamped at –70 mV. Whole-cell voltage clamp recordings were performed using a MultiClamp 700B amplifier, filtered at 8 kHz and digitized at 20 kHz using a Digidata 1550A digitizer (Molecular Devices). Data were acquired and stored using Clampfit 10.4 software (HEKA Electronics) and analyzed with Mini-Analysis (Synaptosoft Inc., Decatur, GA). The neuronal activity from 200.000 hippocampal cells was sampled extracellularly at 10 kHz using MC_Rack software and MEA1060INV-BC system (MultiChannel Systems, Reutlingen, Germany) placed inside of a cell culture incubator in order to provide properly controlled temperature, humidity, and gas composition as described (Bikbaev et al. 2015). The recordings were initiated after a resting period of 30 min after physical translocation of each individual MEAs to the recording system. The off-line analysis was carried out on 600-sec long sessions per MEA at each experimental condition. The detection of spikes was performed after a high-passed (300 Hz) filtering and processing of signals and analyses of neuronal activity were carried out using Spike2 software (Cambridge Electronic Design, Cambridge, UK).

### Statistical analysis

The results are presented as mean ± SEM (standard error of the mean) accompanied by N number of cells in the figure legends. For statistical analysis, Prism 5 software (GraphPad) was used. Mean between two groups was compared with the Student‘s t-test. Means of two or more groups were analyzed by one-way ANOVA (analysis of variance) followed by Tukey’s multiple comparison test for post hoc comparisons. Normalized data were compared using Mann-Whitney test.

## Acknowledgements.

We thank to Dr. Dirk Montag for kindly providing *Nptn^-/-^* mice for breading and to Kathrin Pohlmann for her technical work. S.K.V. thanks to the GRK 1167. A.M. thanks the federal state Saxony-Anhalt and the European Structural and Investment Funds (ESF, 2014-2020), project number ZS/2016/08/80645. R.A.M. thanks the FONDECYT Grant No. 1181260. M.P. thanks to Leibniz Association (SheLi J28/2017). R.H-M. thanks to Center for Behavioral and Brain Sciences (LSA-fellowship) and DAAD project number 57514679. R.H-M., C.I.S., and E.D.G. thanks to SFB 854 and BMBF 01DN17002.

## Author Contributions

S.K.V. and A.M. conducted most of the experiments and raw data analysis. L.J. conducted PMCA experiments. A-C.L. and R.R. made and characterized constructs. J.H. conducted SPR experiments. R.A.M. conducted in silico modelling. M.P. conducted Patch Clamp experiments. R.H.M. supported experiments, conducted MEA and Patch Clamp experiments, and wrote manuscript draft. R.H.M, M.K. M.P., M.N., C.I.S., and E.D.G. data interpretation. All authors contributed to the manuscript final version.

---

**Figure S1(related to Figure 2).**
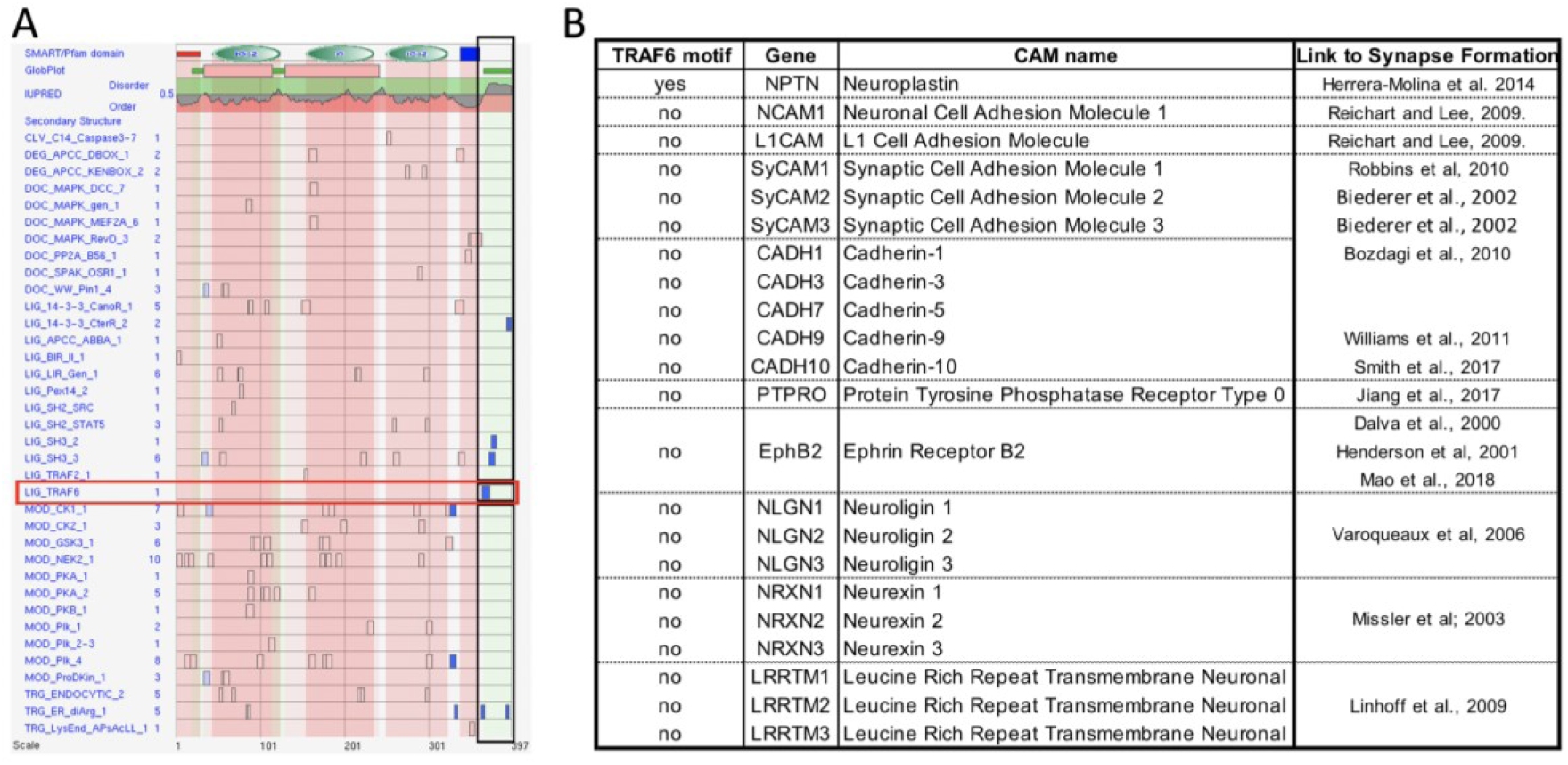
**A.** Data base-based identification of an intracellular TRAF6 binding site in neuroplastin tail. ELM database (http://elm.eu.org/) read out table showing identified binding motifs in all the mouse neuroplastin structure (top drawing). The three extracellular Ig-like domains are in green, transmembrane in blue, and intracellular tail is not colored. Note that the a single TRAF6 binding motif is identified in the cytoplasmic tail (blue square into the red frame). **B.** The table shows ELM database-based motif analysis for the listed synaptogenic cell adhesion molecules. Note that only neuroplastin display TRAF6 binding motif.

**Figure S2(related to Figure 2).**
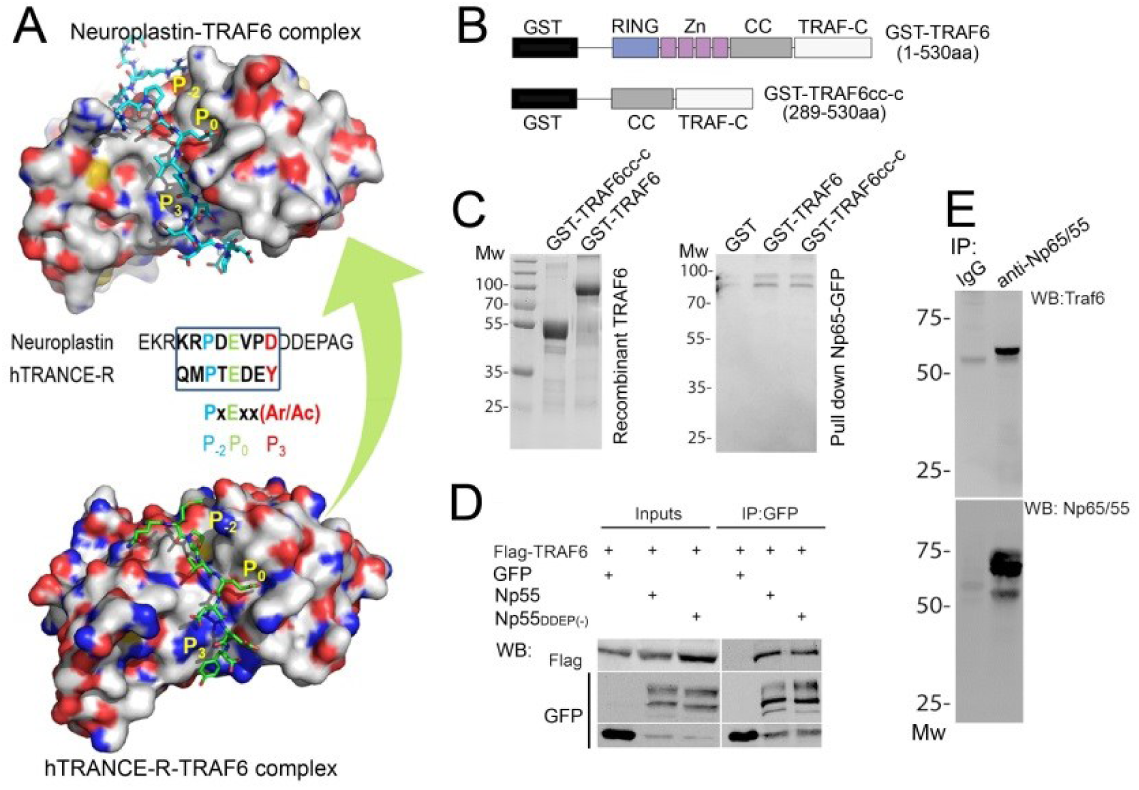
**A.** Modeling neuroplastin-TRAF6 binding. This model is based on the hTRANCE-R-TRAF6 interaction according to provided structures (Ye et al., 2002). The local peptide docking of the tail of neuroplastin (cyan; PBD: **1LB5_A**) is shown on the top and the template complex is shown below (hTRANCE-R in green; PDB: **1LB5_B**). Protein structure similarity (TM-score) = 0.991, Interaction similarity = 108.0, and Estimated accuracy = 0.868. Positions P_-2_, P_0_ and P_3_ in the TRAF6 motif are indicated in both peptide-protein complexes. Protein surfaces are colored based on element (C in white; O in red; N in blue; S in orange). **B.** Representation of GST-TRAF6 and GST-coiled coil-TRAF-C domain (TRAF6_cc-c_) recombinant proteins. RING domain, zinc fingers (Zn), coiled-coil region (CC) and C-terminal domain (TRAF-C) are indicated. **C.** Left: Coomassie gel showing recombinant GST-TRAF6 and GST-TRAF6_cc-c_ recombinant proteins. Right: Western blot showing that both recombinant proteins are pulled-down by Np65-GFP from total extracts of transfected HEK cells. **D**, **E**. Co-precipitation of TRAF6 with neuroplastins from brain and HEK cells homogenates. (**D**) The two Np55 isoforms (with and without DDEP insert) are effective to co-precipitate TRAF6. HEK cells were transfected with the indicated constructs, left to express the tagged proteins for 24 hours, and lysed with RIPA lysis buffer. The extracts were immunoprecipitated with anti-GFP antibody coupled to magnetic beads. Precipitated complexes were resolved by SDS-PAGE and immunoblotted with anti-Flag or anti-GFP antibodies. (**E**) Three-weeks old rat brains were homogenized and lysed with RIPA lysis buffer and incubated with an antibody recognizing all neuroplastin isoforms raised in rabbit (1μg/ml, Smalla et al., 2000) for 24 hours at 4°C. Precipitated proteins were resolved by SDS-PAGE and immunoblotted with pan anti-Np65/55 antibody from sheep or anti-TRAF6 antibody from rabbit.

**Figure S3(related to Figure 3).**
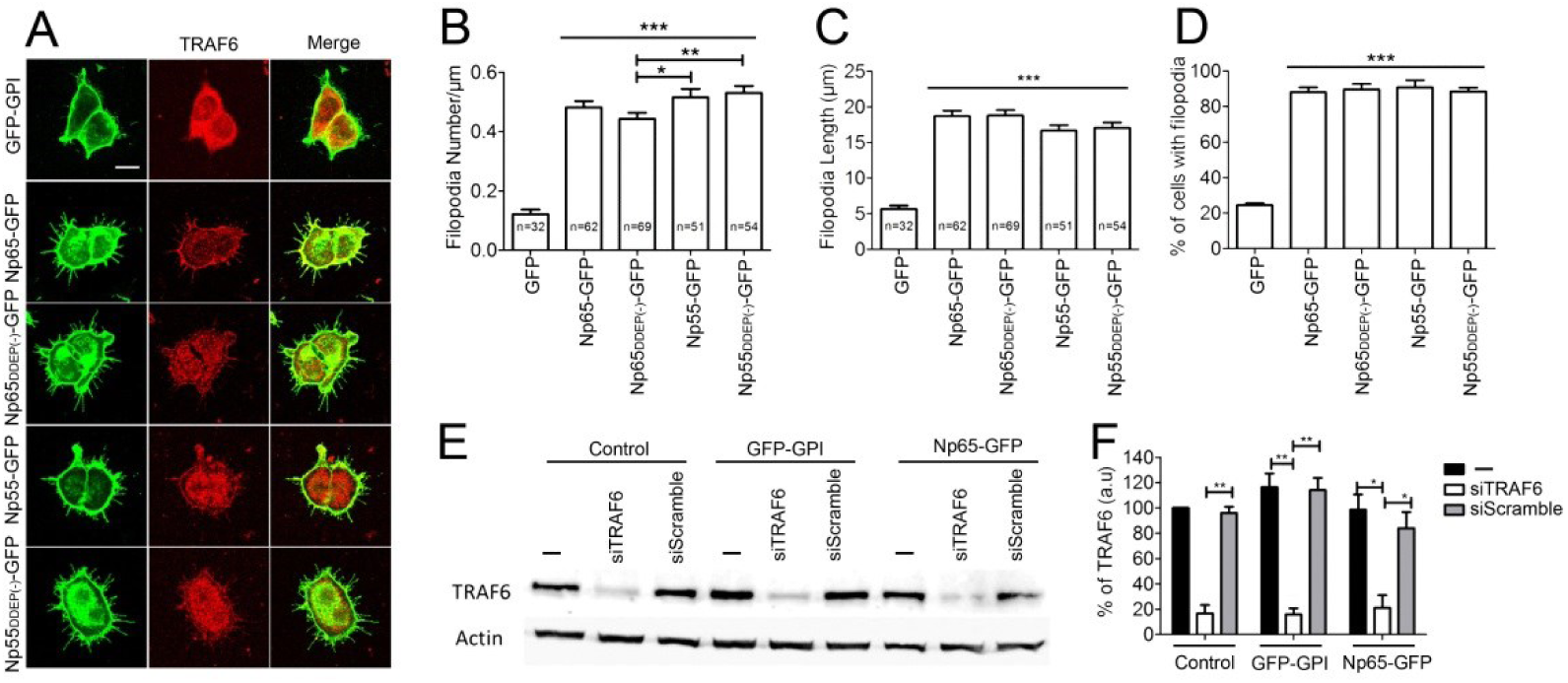
**A**-**D**. The four isoforms of neuroplastin are equally robust to promote translocation of endogenous TRAF6 to the cell membrane and to increase both number and length of filopodia. (**A**) Confocal images displaying representative examples of HEK cell transfected with different neuroplastin constructs and stained for TRAF6 as for Figure 3D. Scale bar=10 µm. (**B**) Number of filopodia (GFP=0.12 ± 0.01 N=32; Np65-GFP=0.48 ± 0.02 N=62; Np65_DDEP(-)_-GFP=0.44 ± 0.02 N=69; Np55-GFP=0.52 ± 0.03 N=51; Np55_DDEP(-)_-GFP=0.53 ± 0.02 N=54) (**C**) Filopodia length (GFP=5.66 ± 0.49; Np65-GFP=18.72 ± 0.77; Np65_DDEP(-)_-GFP=18.79 ± 0.76; Np55-GFP=16.68 ± 0.76; Np55_DDEP(-)_-GFP=17.08 ± 0.72) and (**D**) Percentage of cells with filopodia are displayed as mean ± SEM. Student‘s t-test (**B**,**C**) or with Mann-Whitney test (**D**) were applied. *p<0.05, **p<0.01, and ***p<0.001 vs GFP. **E**, **F**. Assessment of TRAF6 knockdown efficiency upon siRNA treatment. (**E**) Total cell homogenates were resolved by SDS-PAGE and immunoblotted for endogenous TRAF6 and actin to control protein loading. (**F**) The graph shows the densitometric quantification of TRAF6 bands from three independent experiments. **p<0.01 or *p<0.05 using Mann-Whitney test.

**Figure S4(related to Figure 4).**
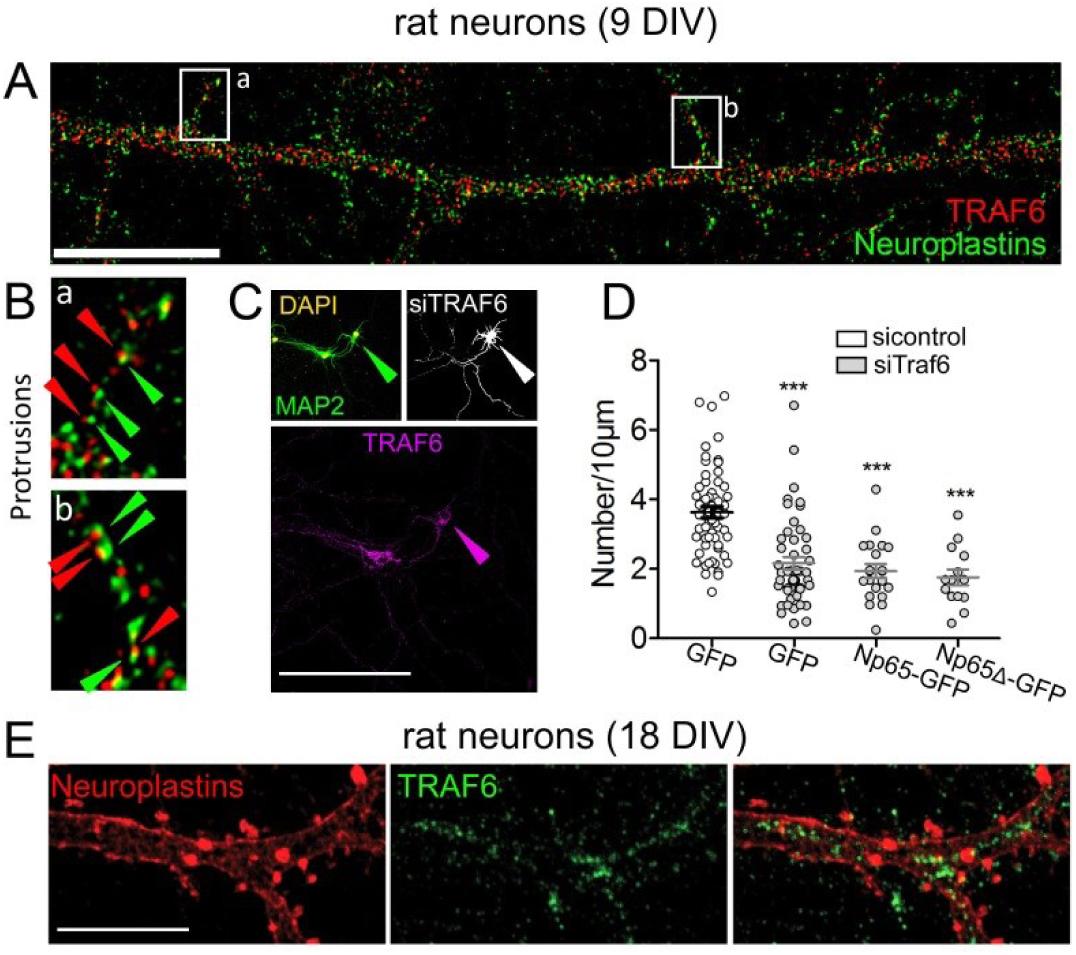
**A**, **B**. Staining of neuroplastin and TRAF6 in methanol-fixed rat young hippocampal neurons. (**A**) Neurons were stained with a pan-antibody recognizing all neuroplastin isoforms and anti-TRAF6 antibody followed by proper fluorophore-tagged secondary antibodies, mounted, and imaged using a 100x objective of a confocal microscope. Scale bar=10 µm (**B**) Digital magnification of dendritic protrusions with co-distributed and co-localized spots of neuroplastin and TRAF6 displayed. For **A** and **B**, images were deconvolved (see methods). **C**, **D**. TRAF6 knockdown counteracts the increase of dendritic protrusions induced by Np65-GFP over-expression in hippocampal neurons. Neurons were co-transfected with either control scrambled siRNA or siRNA against TRAF6 mRNA and with GFP-encoding plasmid (6 DIV). Additionally, neurons were co-transfected with siRNA and Np65-GFP or Np65Δ-GFP. After 72 hours, neurons were stained with anti-MAP2 and anti-TRAF6 antibodies to control neuronal morphology and TRAF6 KD, respectively. Only neurons with ≥60% reduction in TRAF6 immunoreactivity (arrow heads) were considered for the counting of dendritic protrusions. Transfected neurons from four independent cultures were analyzed (si-control GFP=3.73 ± 0.16 N=59; siTRAF6 GFP= 2.16 ± 0.18 N=49; siTRAF6 Np65-GFP= 2.09 ± 0.16 N=22; siTRAF6 Np65Δ-GFP= 1.69 ± 0.17 N=14). ***p<0.001 vs. si-control GFP using Student‘s t-test. Scale bar=100 µm. **E**. Neuroplastin and TRAF6 in mature hippocampal neurons. Staining was performed as in **A**. TRAF6 does not co-localize with neuroplastin in mature neurons. Scale bar=10 µm.

**Figure S5(related to Figure 4).**
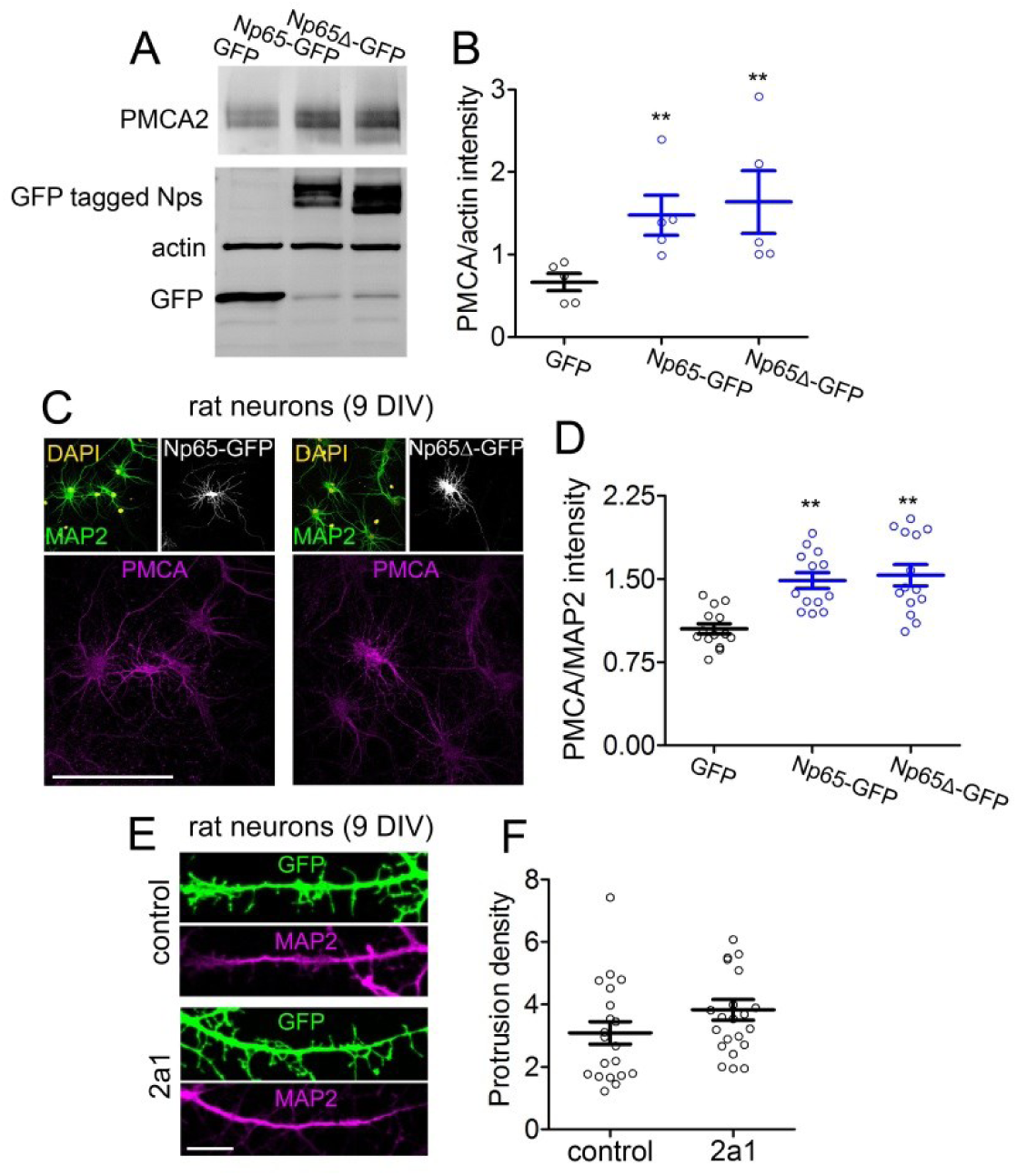
**A**, **B.** Protrusion formation does not depend on Neuroplastin-PMCA interaction. (**A**) Np65-GFP and Np65Δ-GFP equally increase total PMCA2 levels in HEK cells. Cells were transfected with the indicated constructs, harvested 24 hours later and lysed with RIPA lysis buffer. Western blot analysis shows that levels of PMCA2 are increased upon co-transfection with Np65-GFP and with Np65Δ-GFP as indicated. Blotting of actin is used to control loading. (**B**) Quantification of PMCA2 blots are normalized to actin using data from five independent experiments. **p<0.01 vs. GFP using Mann-Whitney test. **C**, **D.** Np65-GFP and Np65Δ-GFP lacking intracellular TRAF6 domain are equally effective to increase PMCA expression in young hippocampal neurons. (**C**) At 7DIV hippocampal neurons were transfected with plasmids encoding Np65-GFP or Np65Δ-GFP. At 9DIV, neurons were fixed and stained with an anti-MAP2 and anti-pan-PMCA antibodies. Scale bar=10 µm. (**D**) Quantification of the intensity of PMCA immunofluorescent signal normalized to MAP2 signal using data from 13-20 neurons per group from three independent cultures. **p<0.01 vs. GFP using Student‘s t-test. (GFP=1.05 ± 0.04 N=14; Np65-GFP=1.48 ± 0.07 N=13; Np65Δ-GFP= 1.53 ± 0.09 N=14). **E**, **F**. Hippocampal neurons (DIV8) were transfected with GFP-expressing plasmid. After 24 hours, neurons were incubated with the PMCA inhibitor Caloxin 2a1, fixed, immunostained with an anti-GFP antibody and an anti-MAP2 antibody followed by proper secondary antibodies, and imaged using a confocal microscope with a 63X objective under 3X digital zoom factor. (**F**) Protrusion density in control and Caloxin 2a1 treated neurons from two independent cultures (control=3.08 ± 0.35 N=20; 2a1= 3.83 ± 0.39 N=25).

